# Mapping disease traits onto spatiotemporal domain landscapes through scalable multi-sample integration of spatial transcriptomics

**DOI:** 10.64898/2025.12.15.694341

**Authors:** Songming Zhang, Shifu Luo, Yuxiao Luo, Shuquan Su, Ling Liu, Ying Shi, Wujun Li, Tian Tian, Jinyan Li

**Affiliations:** School of Computer Science, Nanjing University, Nanjing, China; Faculty of Computer Engineering and Artificial Intelligence, Shenzhen University of Advanced Technology, Shenzhen, China; Shenzhen Institute of Advanced Technology, Chinese Academy of Sciences, Shenzhen, China; School of Computer Science, Centre for Human Brain Health, University of Birmingham, Birmingham, UK; School of Computer Engineering & Science, Shanxi University, Taiyuan, China; School of Artificial Intelligence, Wuhan University, Wuhan, China

**Keywords:** *Spatial transcriptomics*, *integrative analysis*, *hypergraph construction*

## Abstract

High-resolution spatial transcriptomics (ST) is generating massive datasets involving multiple tissue sections, continuous developmental stages, or various disease conditions. Yet, their scale, biological heterogeneity, and substantial batch effects hinder data integration for further analysis, particularly the temporal interpretation of disease-associated spatial domains within tissue organization landscapes. Here, we present STUltra, a scalable hypergraph framework that integrates multi-sample ST data for precise domain detection and genomewide association study (GWAS)-based disease trait mapping. From ST datasets, STUltra first constructs interval-sampled integrative hypergraphs, in which hyperedges capture tissue neighborhoods within the slices as well as shared biological contexts across the slices. It then combines a robust graph autoencoder with contrastive learning to learn batch-corrected, spatially informed embeddings for identifying spatial domains across the tissue sections. These embeddings are then coupled with GWAS statistics to map trait-associated signals onto the integrated tissue sub-structure landscapes spanning different sections, developmental stages, and disease conditions. STUltra substantially outperforms competing methods, especially being much faster, on diverse integration benchmarks, enabling efficient processing of datasets containing millions of spots. As a case study on a mouse embryo dataset, STUltra recovered continuous developmental programs and mapped congenital heart disease signals beyond cardiac regions to broader programs including vascular and mesenchymal domains. On a mouse Alzheimer’s disease dataset, STUltra prioritized localized AD-associated genetic signals to microglia and a disease-expanded *C1QA*/*HEXB*-high astrocyte state. Together, STUltra provides a scalable framework to bridge the gap between spatiotemporal tissue structures and complex disease traits, facilitating discoveries from previously unrecognized molecular and cellular architectures across diverse contexts.

## 1. Main

Cellular grouping and neighborhood interactions evolve spatiotemporally, underlying gradual tissue development and disease progression. Spatial transcriptomics (ST) technologies facilitate the developmental studies of tissue structures by measuring gene expression in its native tissue context. Recent advances in high-resolution and imaging-based ST technologies have expanded spatial profiling to millions of spots per tissue section (*e.g.*, Visium [1], Xenium [2] Visium HD [3], and Stereo-seq [4]), and thus providing unprecedented potential to uncover novel insights into cellular diversity and functional programs within the structured tissue landscapes.

One of the primary tasks in ST data analysis is to accurately identify distinct sub-structures or domains from these tissue sections [5–7]. With the increasingly accumulated ST datasets across tissue samples, sequencing platforms, developmental stages, and disease conditions, the analysis is currently shifting from single-section characterization toward large-scale multi-sample integration. Such spatiotemporal integration will enable harmonization of datasets generated by different technologies, identification of shared and context-specific tissue architectures, comparison of spatial domains across biological conditions, and subsequent reconstruction of spatiotemporal development programs that cannot be fully resolved from any single section [8]. However, existing methods often rely on representations designed for individual sections or pairwise alignment, which ignore both spatial continuity and local variability (*e.g.*, GraphST [9], SpaVAE [10], HERGAST [11]). Therefore, scalable integration methods that jointly model spatial organization and biological variation are needed for spatiotemporal domain identification in ultra-large multi-sample ST datasets.

Several computational methods have been developed for multi-sample ST alignment and integration. However, these tools were developed and evaluated on datasets containing 10,000–100,000 cells, leaving their scalability to ultra-large multi-sample datasets uncertain. PASTE [12] and 3D-OT [13] align adjacent tissue sections to infer spatial continuity, while STitch3D [14] and SEDR [15] construct spatial neighbor graphs for integration and spatial domain identification. These methods are effective for related or adjacent sections, but their reliance on pairwise alignment or predefined spatial correspondence may limit their use in heterogeneous datasets collected. BANKSY [16] builds joint spatial representations by combining expression and neighborhood information, whereas STAligner [17] and CellCharter [18] use graph neural networks to model gene expression and spatial context across sections. Although these approaches improve spatial integration, their steps for graph construction and model training become increasingly expensive when the number of spatial spots grows [19, 20]. In addition, aggressive batch correction or joint embedding may blur condition-associated local domains, whereas insufficient correction can retain section-specific technical variation [21].

Beyond identifying distinct tissue structures, another important task is to determine how these spatially organized sub-structures are related to the genetic basis of human complex traits. Recent methods, such as gsMap [22], have combined ST data with genome-wide association study (GWAS) statistics to map trait-associated cells within individual tissue sections to demonstrate the value of spatially resolved genetic association analysis. However, many complex traits and diseases emerge and progress across several developmental stages, anatomical regions, and pathological conditions, and their relevant spatial contexts may not be fully captured by a single section [22–24]. Extending trait mapping from individual sections to integrated multi-section and time-resolved tissue architecture landscapes may therefore provide a broader view of how disease signals are distributed across spatial and biological contexts.

Here, we introduce STUltra, a scalable hypergraph framework for integrating ultra-large multi-sample ST datasets and subsequently used to map domain-associated traits. From ST datasets, STUltra first constructs interval-sampled integrative hypergraphs to model intra-slice tissue neighborhoods and cross-slice biological correspondences in a computationally efficient manner. Specifically, intraslice hyperedges connect sampled spatial spots with their local spatial neighbors, whereas interslice hyperedges link spots with similar gene expression profiles and local neighborhood contexts across the sections. This design captures both local tissue organization and cross-section biological correspondences while reducing the computational cost of large-scale integration. A robust graph autoencoder combined with contrastive learning then generates batch-corrected, spatially informed embeddings for identifying integrated spatial domains to create a spatiotemporal tissue landscape. These integrated representations are further coupled with GWAS statistics to map association-level trait signals onto the tissue architecture landscapes across the sections, developmental stages, and disease conditions.

We evaluated STUltra on diverse biological settings, including human cortical sections, mouse brain sections profiled at different orientations and platforms, colorectal cancer sections, whole mouse embryo sections covering organogenesis, and disease-associated tissue states. Across these settings, STUltra improved spatial domain integration performance, aligned commonly shared biological structures across the slices and platforms, well preserved local tissue architecture in these ultra-large high-resolution data; and its performance can scale to more than 1,000,000 spatial spots on a single 24-GB GPU. Furthermore, combining GWAS statistics with STUltra representations has enabled effective prioritization of candidate trait-associated spatial domains across various developmental and disease contexts. For example, we highlighted congenital heart disease-associated developmental contexts in mouse embryos and reactive glial contexts associated with Alzheimer’s disease. Overall, STUltra provides a scalable framework for ultra-large ST integration and is effectively applicable for prioritizing trait-associated tissue structures across diverse spatial contexts.

## 2. Results

### 2.1. Overview of STUltra

STUltra integrates spatial transcriptomic data from multiple ultra-large ST slices collected under various biological conditions mainly through our three computational components (Fig. 1): (i) interval-sampled hypergraph construction to jointly encode intra-slice and inter-slice neighborhoods while ensuring scalability for high-resolution slices; (ii) intra-slice representation learning via a robust graph autoencoder; (iii) biologically coherent integration across slices through contrastive learning.

**Figure 1.**
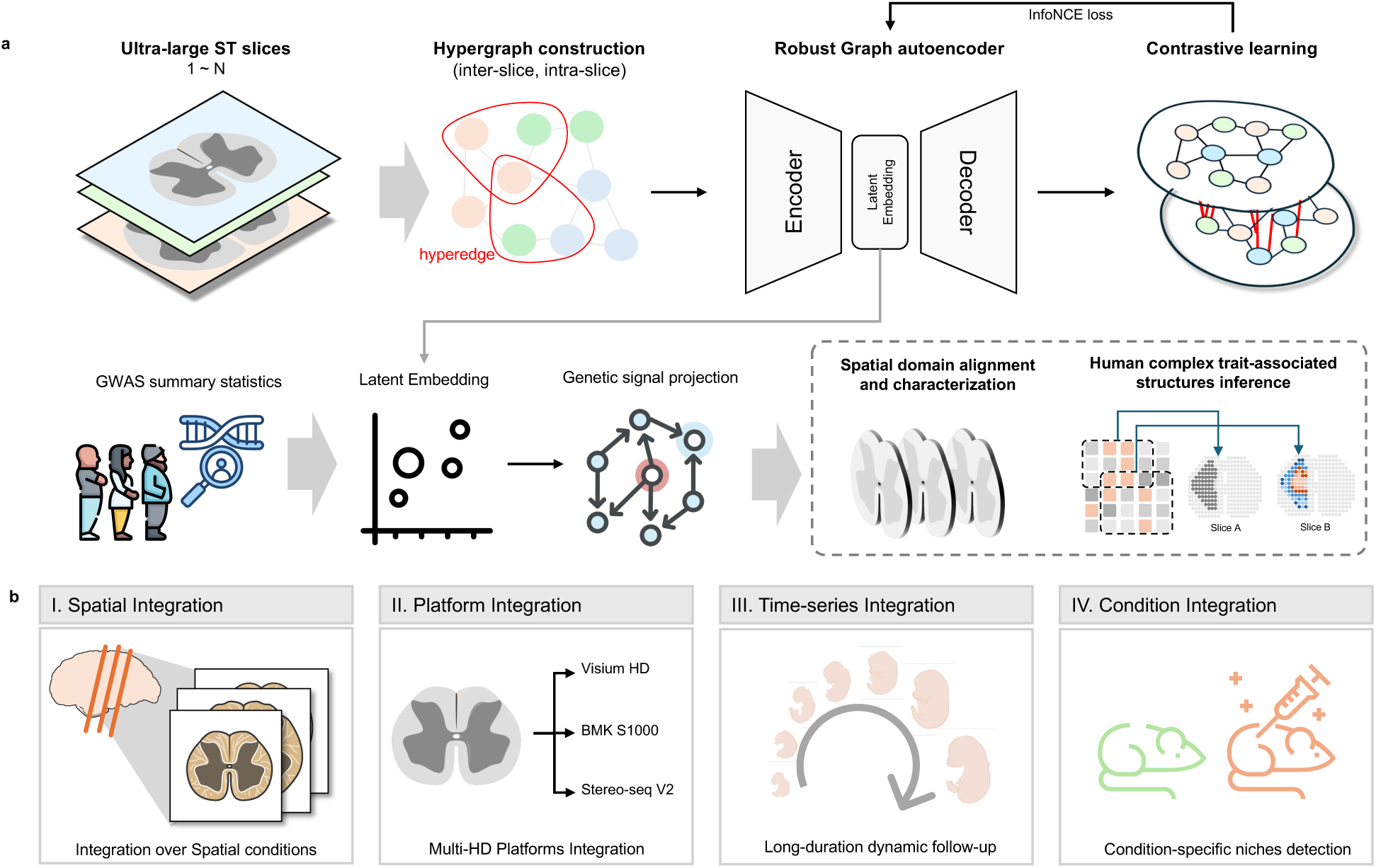
Overview on STUltra. (a) Workflow for STUltra. STUltra first performs interval-sampled integrative hypergraph construction from multi-slice ST data, where spatial input units and their gene expression profiles are organized through both intra-slice and inter-slice correspondence hyperedges with interval-sampled subsets. STUltra further employs a robust graph autoencoder and contrastive learning under the InfoNCE loss to learn spatially consistent embeddings until batch-corrected embeddings are generated. Leveraging the integrated representations learned by STUltra, GWAS summary statistics can be mapped onto ST slices from diverse sources, enabling the identification of trait-associated cells and tissue structures. (b) STUltra enables the integration of high-definition ST slices collected from different biological settings, including different locations in spatial regions, different sequencing platforms, different slice-section time points, or different experimental conditions.

In detail, STUltra first constructs an integrative hypergraph from multi-slice ST data, where intraslice neighborhood hyperedges preserve spatial adjacency within each tissue section, and inter-slice neighborhood hyperedges link biologically corresponding local contexts across the slices. To handle the extremely large number of spatial input units in each high-resolution slice, we carry out an interval gridding strategy to sub-layer and reorganize the slice as patches. Next, the high-order neighborhood relations are projected onto sparse graph representations, and then STUltra employs a robust graph autoencoder to learn low-dimensional embeddings. These embeddings are then iteratively aligned, intersecting the slices through contrastive learning with the InfoNCE loss.

In the contrastive alignment process, positive pairs are constructed from spatial nodes that share similar transcriptional signatures across the slices, while negative pairs are sampled within the same slice from nodes that are both spatially distant and transcriptionally dissimilar. Importantly, STUltra does not rely on spatial distance alone; negative sampling is jointly constrained by the loss function, which reduces the risk of incorrectly separating biologically similar regions with shared molecular patterns. The InfoNCE objective encourages embeddings of biologically similar regions to be close while preventing the integration process from being dominated by batch-specific shifts.

Furthermore, leveraging these integrated representations, GWAS summary statistics are projected onto spatially resolved tissue landscapes for the purpose of identifying trait-associated domain cells and structures. In detail, local expression patterns from these integrated spatial domains are first summarized as gene-level spatial scores, which are then assigned to SNP annotations through SNP-gene mapping. These spatially informed SNP annotations are then tested for a trait heritability enrichment combined at the regional level using the Cauchy combination test of point-wise association *P* values, yielding interpretable maps of the genetic association spatiotemporally across the multiple ST slices of temporal continuity.

### 2.2. Precise integration for slices under different spatial configurations

To evaluate the performance of STUltra in delineating tissue regions in spatial context, we tested the method under various spatial configurations, such as vertical sections, horizontal sections, and high-resolution imaging modalities. The human dorsolateral prefrontal cortex (DLPFC) dataset represents a homogeneous cortical structure, while the mouse brain dataset captures a heterogeneous composition across multiple anatomical regions. On these datasets, we benchmarked STUltra against recent integration methods, including Harmony, STAligner, STitch3D, SEDR, SpaGIC, and SpaBatch, and quantitatively evaluated the performance using the Adjusted Rand Index [21] (ARI) and Normalized Mutual Information (NMI) on manually annotated datasets to highlight STUltra’s superior performance.

Overall, STUltra achieved the highest scores, with an average score 0.63 for ARI and an average score 0.72 for NMI over all the slices (Fig. 2b). In particular, for slice #151673, STUltra reached an ARI 0.67 and NMI 0.78, clearly separating the cortical layers (Layer 1–6) and the white matter (WM) in both the UMAP and spatial domain visualizations. Moreover, the identified cell-type proportions within each layer were closely aligned with the manual annotations.

**Figure 2.**
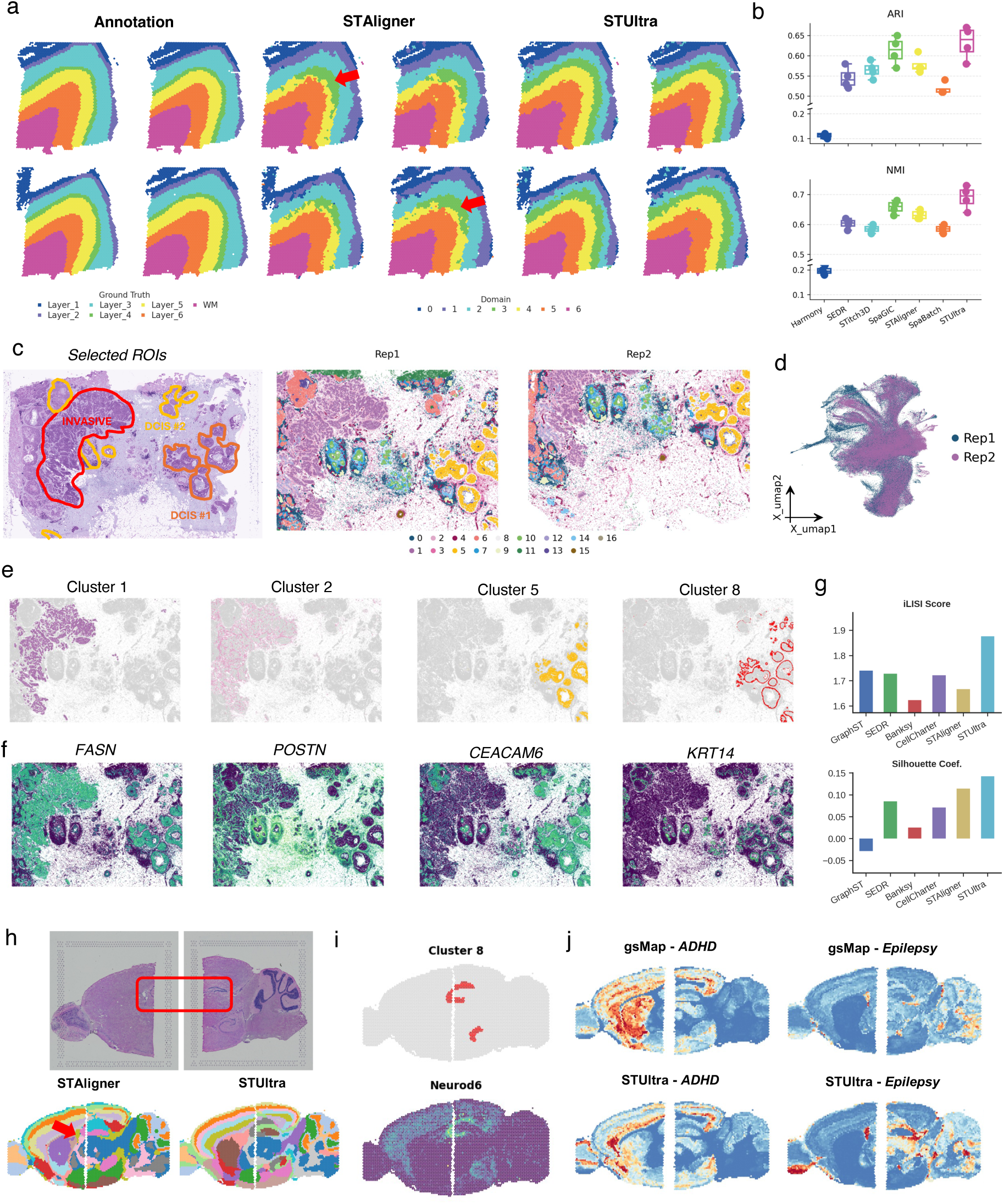
STUltra enables better integration under diverse spatial contexts. (a) Manual annotations, STAligner, and STUltra results derived from four slices in the DLPFC dataset. (b) ARI and NMI scores for seven integration methods: Harmony, STAligner, STitch3D, SEDR, SpaGIC, SpaBatch, and STUltra. (c) H&E image and spatial domains derived by STUltra from two slices of the human breast cancer (BC) 10x Xenium dataset. (d) UMAP visualization colored by slice for the human BC 10x Xenium dataset. (e) Spatial visualization of tumor clusters derived by STUltra from slice Rep1 of the human BC 10x Xenium dataset. (f) Spatial expressions of marker genes in the BC-related regions. (Clusters 1, 2, 5, and 8). (g) iLISI and Silhouette scores for six integration methods: GraphST, SEDR, Banksy, CellCharter, STAligner, and STUltra. (h) The H&E-stained image of the mouse brain anterior and posterior slices, their corresponding specific domains, and spatial domains identified by STAligner and STUltra. (i) Cluster 8 in the mouse brain (top) and its corresponding gene marker *Neurod6* (bottom) returned by STUltra. (j) Trait-mapping results returned by gsMap and STUltra for ADHD and Epilepsy with colours indicating the significance of association.

However, the existing methods exhibited inferior layer resolution: Harmony produced boundaries with high noise and separated only the white matter structure correctly; SpaGIC and STitch3D suffered from over-correction, leading to disordered Layer 6 allocation; and STAligner showed reduced smoothness in the identification of Layer 4 with a smaller proportion of matching cells. On the other hand, STUltra accurately reconstructed spatial trajectories that are topologically consistent with the layered organization of the cortex, exhibiting smooth transitions from the layer surface to WM (Supplementary Fig. S7). The inferred cross-slice-sharing domains exhibited coherent spatial continuity and could be validated by the enriched expression of the canonical layer-specific marker genes such as HPCAL1 for Layer 1, PCP4 for Layer 4, and MBP for the WM domain (Supplementary Fig. S6).

We also applied STUltra to a high-resolution ST dataset generated by 10x Xenium from two adjacent human breast cancer tissue sections [25]. Breast cancer is highly heterogeneous, exhibiting substantial intra- and inter-tumoral variation in both histological and molecular features. From this dataset with two slices, STUltra successfully identified and separated multiple tumor domains (Figure 2c-f, Supplementary Figure S15), including ductal carcinoma in situ (DCIS; clusters 5 and 8) representing non-invasive lesions, and invasive ductal carcinoma (IDC; clusters 1 and 2), as well as several tumor microenvironment (TME)-related domains such as immune-associated regions (cluster 0), tumor stroma (cluster 7), and the myoepithelial layer (domain 3). These spatial domains detected via STUltra exhibited spatial smoothness and compactness, while the enrichment of established marker genes further validated their biological significance, and they were also consistent with ROIs provided by pathologists (Figure 2e-f). For example, FASN2 [25], an invasive tumor marker, was highly enriched in IDC cluster 1. CEACAM6, a cell adhesion molecule whose expression is negatively correlated with cell differentiation, showed strong enrichment in DCIS clusters 5 and 8 [25, 26]. Quantitative evaluation using iLISI and Silhouette scores further showed that STUltra achieved a more favourable balance between cross-slice mixing and preservation of spatial-domain structure in comparison with GraphST, SEDR, Banksy, CellCharter, and STAligner (Fig. 2g).

To further examine the capability of STUltra to integrate heterogeneous tissue slices, we used two sagittal sections from a mouse brain representing the anterior and posterior regions. Of these integration methods, only STUltra and STAligner successfully identified the commonly shared clusters corresponding to the cortex, hippocampus, and thalamus (Fig. 2h). Moreover, only STUltra accurately reconstructed the trisynaptic circuit in the ventral hippocampus and revealed smoother laminar patterns within the cortex, with the spatial domains further confirmed by the expression of the representative marker gene *Neurod6* (Fig. 2i).

To evaluate whether STUltra preserves anatomical structures and improves trait-mapping resolution after our integration, we compared STUltra with gsMap on the ADHD and epilepsy GWAS signals in multi-integrated mouse brain slices [22]. For ADHD, STUltra revealed continuous and significant spatial enrichments spreading across key brain regions, including the cortex and hippocampus regions, with – log_10_(*P*) values reaching approximately 2.5–3.0 along the anatomical boundaries; by contrast, gsMap produced weaker and more diffuse signals, which were much less helpful for distinguishing specific brain regions. For epilepsy, STUltra identified multiple focal hotspots in the forebrain, hippocampus, and subcortical structures, consistent with seizure-related neural circuits, whereas gsMap remained close to the background level under the same colour scale with only a few scattered weak signals. These results suggest that STUltra has better preserved inter-slice anatomical structures after our integration, enabling more precise localization and assessment of the genetic risk for the complex psychiatric and neurological traits than gsMap did (Fig. 2j).

We also validated our STUltra’s spatiotemporal integration capabilities. The experiments were conducted on consecutive coronal slices from the Mouse Hypothalamus dataset [27], which comprises of five slices generated using the imaging-based MERFISH platform. In this dataset, eight anatomical regions were manually annotated, namely the bed nucleus of the stria terminalis (BST), medial preoptic area (MPA), medial preoptic nucleus (MPN), paraventricular nucleus (PV), paraventricular hypothalamic nucleus (PVH), paraventricular thalamic nucleus (PVT), third ventricle (V3), and fornix (fx). We compared STUltra’s performance with two benchmark methods: STAligner and SEDR (Supplementary Figure S2a). SEDR’s results exhibited more dispersed domain boundaries with a lower ARI score. In comparison, STUltra outperformed these two deep learning methods in spatial domain identification, achieving the highest ARI mean score of 0.449, representing a 10% improvement. Take V3 as an example, which is a midline brain structure filled with cerebrospinal fluid and is located within the diencephalon, primarily between the thalamus and hypothalamus. STUltra accurately detected this domain, drawing distinguishable boundaries from the surrounding regions. We further conducted differential expression analysis among multiple spatial domains (results visualized via dot plots, Supplementary Figure S2c); this analysis identified *Cd24a* and *Nnat* as marker genes. Additionally, we made a visualization of the marker gene *Cd24a* for the five sections in 3D space (Supplementary Figure S2b). Overall, STUltra is capable of well aligning the related clusters among the slices under varying spatial contexts. More ablation analysis results on STUltra modules are shown in Supplementary Appendix S1. All experiments were independently repeated three times, and the reported results correspond to the average performance over all the runs.

### 2.3. Identification of tumor-adjacent local spatial domains in colorectal cancer slices

With high mortality and poor prognosis, colorectal cancer (CRC) remains a major global health burden, underscoring the need for improved early detection and prognostic biomarker discoveries. ST can map gene expression onto the space and reveal complex cellular responses that are not visible by classic histology. We applied STUltra to integrate spatial transcriptomics data of two 10x Genomics Visium HD colorectal cancer slices (>1.2 million bins, profiling >10,000 genes per bin) to demonstrate its scalability and utility for novel biological discovery, as other methods failed due to a high computational cost.

STUltra identified 11 clusters and provided a smooth yet detailed delineation of tumor stroma regions (Fig. 3b, Supplementary Fig. S12). In contrast, BANKSY failed to consistently align the tumor-associated local spatial structures in the colorectal cancer dataset; it could not even properly match the commonly shared spatial domains between the two adjacent tissue sections, leading to fragmented and inconsistent cross-section spatial organization. To further investigate its immune dynamics, we extracted immune cell populations located near the tumor cells (Cluster 0, Supplementary Fig. S9). Within a close spatial radius, macrophage-enriched domains were spatially associated with tumor regions. In fact, for this dataset, macrophages were distributed variably around the tumor cells, forming clusters, strands, or gland-like structures surrounding epithelial-like regions, posing challenges for accurate integration. Notably, STUltra precisely aligned these intratumoral macrophages at these two slices, highlighting its advantage in resolving complex spatial interactions and in uncovering heterogeneity within tumor regions.

**Figure 3.**
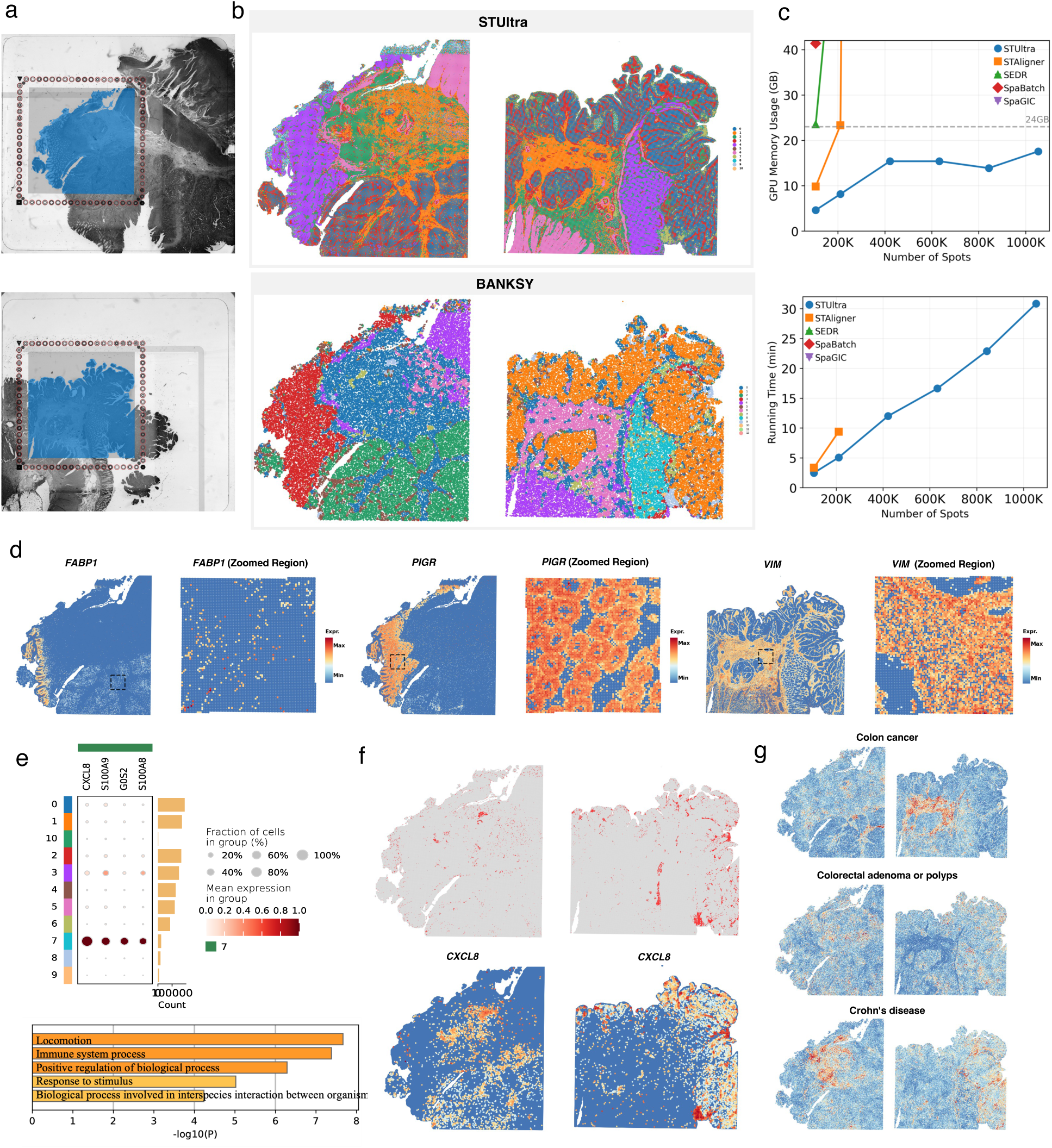
STUltra achieved precise identification of intra-tumoral local spatial domains in two human colorectal cancer slices. (a) The H&E-stained images from Visium HD human colorectal cancer slices (8*μm*). (b) Visualization of the aligned spatial domains identified by STUltra and BANKSY. (c) Runtime cost and GPU memory usage by STUltra. Upper: GPU memory; Bottom: Runtime cost. All experiments were performed on a server with an Intel Xeon(R) Platinum 8360Y CPU and an NVIDIA GeForce RTX 4090 GPU (24 GB memory). (d) Corroboration of selected gene markers (FABP1, PIGR, and VIM) with their spatial localizations. For each sample, the tissue-level view is shown on the left with the inset region outlined by a black box, and the corresponding zoom-in view is shown on the right. (e) Dot plot of the highly expressed genes in Cluster 7 identified by STUltra. (f) Visualization of spatial domains for Cluster 7 (top). The significantly enriched biological pathways, as determined using the Metascape web tool (bottom). (g) Trait-mapping results for Colon cancer, Colorectal adenoma or polyps, and Crohn’s disease with colours indicating the significance of association.

We quantified spatial relationships by computing cluster co-occurrence probabilities (Supplementary Fig. S9). The co-occurrence profile for the tumor domain showed the highest association with the tumor margin, which also reinforces the interpretability of our annotations. We then visualized marker genes in the regions of interest, including FABP1 for Cluster 0 (tumor), PIGR for Cluster 4 (goblet cells and enterocytes), and VIM for Cluster 3 (fibroblasts); These markers show strong spatial co-expression and match with previously reported patterns (Fig. 3d). Around a calcified region, STUltra detected a distinct and sparsely expressed domain. (Cluster 7; Fig. 3e, Supplementary Fig. S14). To characterize this domain, we summarized its highly expressed genes into a metagene, which exhibited a clear and localized spatial pattern corresponding to the cluster. Pathway enrichment analysis of this metagene revealed significant up-regulation of immune-related processes, including antigen presentation, cytokine signaling, and lymphocyte activation, consistent with the observed local aggregation and infiltration of immune cells. These findings suggest that this novel domain represents an immune-enriched tumor-associated spatial domain (Fig. 3f).

We also mapped trait associations onto the integrated CRC tissue landscape to examine whether genetic risk signals were localized to distinct spatial microenvironments (Fig. 3g). Within the Visium HD colorectal cancer tissue, distinct gastrointestinal traits exhibited divergent spatial association patterns. Colon cancer signals were preferentially localized to smooth muscle-associated structures, whereas colorectal adenoma and polyp signals were enriched in endothelial–myeloid regions, consistent with the involvement of stromal remodeling and vascular-associated microenvironments during early lesion development. In contrast, Crohn’s disease displayed the strongest associations with myeloid-enriched structures, reflecting its close relationship with immune-driven inflammatory processes. These trait-specific spatial distributions suggest that genetically associated signals are organized within distinct tissue architectures and disease-relevant microenvironmental contexts. Together, these results highlight the ability of our framework to infer trait-associated structures from integrated ultra-large spatial transcriptomic landscapes and connect genetic risk to spatially resolved tissue organization.

With recent improvements in ST resolution, each tissue slice can now contain hundreds of thousands of spatially resolved input observations that cover an extensive field of view. Such ultra-large slices impose substantial computational demands, requiring integration methods that should be both memory- and time-efficient to run on personal servers. To assess the scalability of STUltra, we measured its runtime cost and GPU memory usage on the CRC datasets when a personal server equipped with a 24 GB GPU was used (Fig. 3c). For integrating this dataset containing more than 1,000,000 input units, STUltra completed the integration in 31.5 minutes, while our sampling strategy allows further control over peak GPU memory consumption. In contrast, existing deep learning-based tools such as SEDR and SpaGIC were limited to fewer than 100,000 units on the same hardware, highlighting the markedly higher efficiency of STUltra for large-scale integration.

### 2.4. Enhanced integration for slices from different high-resolution platforms

To demonstrate STUltra’s cross-platform integration capability, we used three adult mouse brain datasets [19] generated by different high-resolution platforms, including Visium HD, BMK S1000, and Stereo-seq [4], which together contain over 1.2 million spatial input units (Fig. 5a). The Visium HD data were produced with 10x Genomics’ enhanced Visium assay for high-definition whole-slide sections, the BMK S1000 data were generated on a bead-patterned array that achieves subcellular resolution, and Stereo-seq used high-density barcoded chips to provide nanoscale, large-area coverage. In addition, due to differences in experimental setups, data generated by different platforms often exhibit substantial heterogeneity, such as variations in spatial resolution, capture efficiency, and coverage of local structures, which further complicates cross-platform integration.

**Figure 4.**
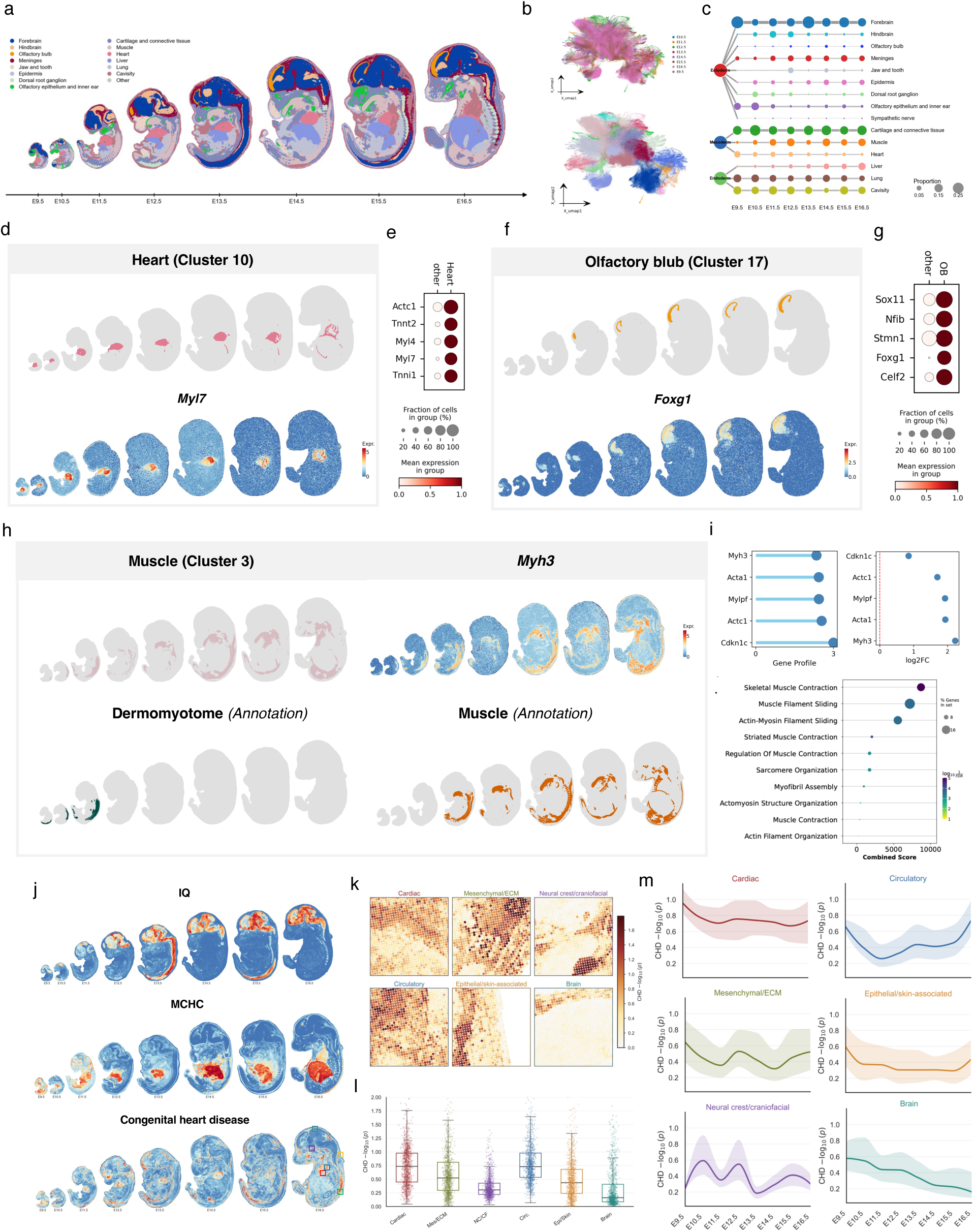
STUltra uncovered spatiotemporal development dynamics inside long-term Stereo-seq embryo data. (a) Time-series Stereo-seq data of mouse embryo development from E9.5 to E16.5 with the annotations. (b) UMAP visualization of spot embeddings, with colors indicating section index and spatial domain labels as determined by STUltra. (c) Directed acyclic graph showing the developmental dynamics of tissue domains. (d,e) Spatial domains (top) related to the heart and corresponding marker gene. expression (bottom). (f,g) Spatial domains (top) for the olfactory bulb and their marker genes (bottom). (h,i) Comparison of STUltra-integrated results (top) and the original annotation (bottom). STUltra unifies reference dermomyotome and muscle labels into a myogenic domain at multiple stages; *Myh3* expression is enriched at later stages within this domain, consistent with late-stage muscle differentiation. Gene Ontology (GO) analysis is shown for factor-specific genes in (h). (j) Trait-mapping results associating spatial domains with IQ, MCHC, and congenital heart disease (CHD), with color representing the significance of association. (k) Representative high-resolution view of CHD-associated regions highlighted from (j). (l) Box plots showing the CHD association significance for each identified spatial region at the E16.5 slice. (m) Temporal dynamics of CHD-association scores for exemplar regions from E9.5 to E16.5, demonstrating how trait associations evolved during embryo development.

**Figure 5.**
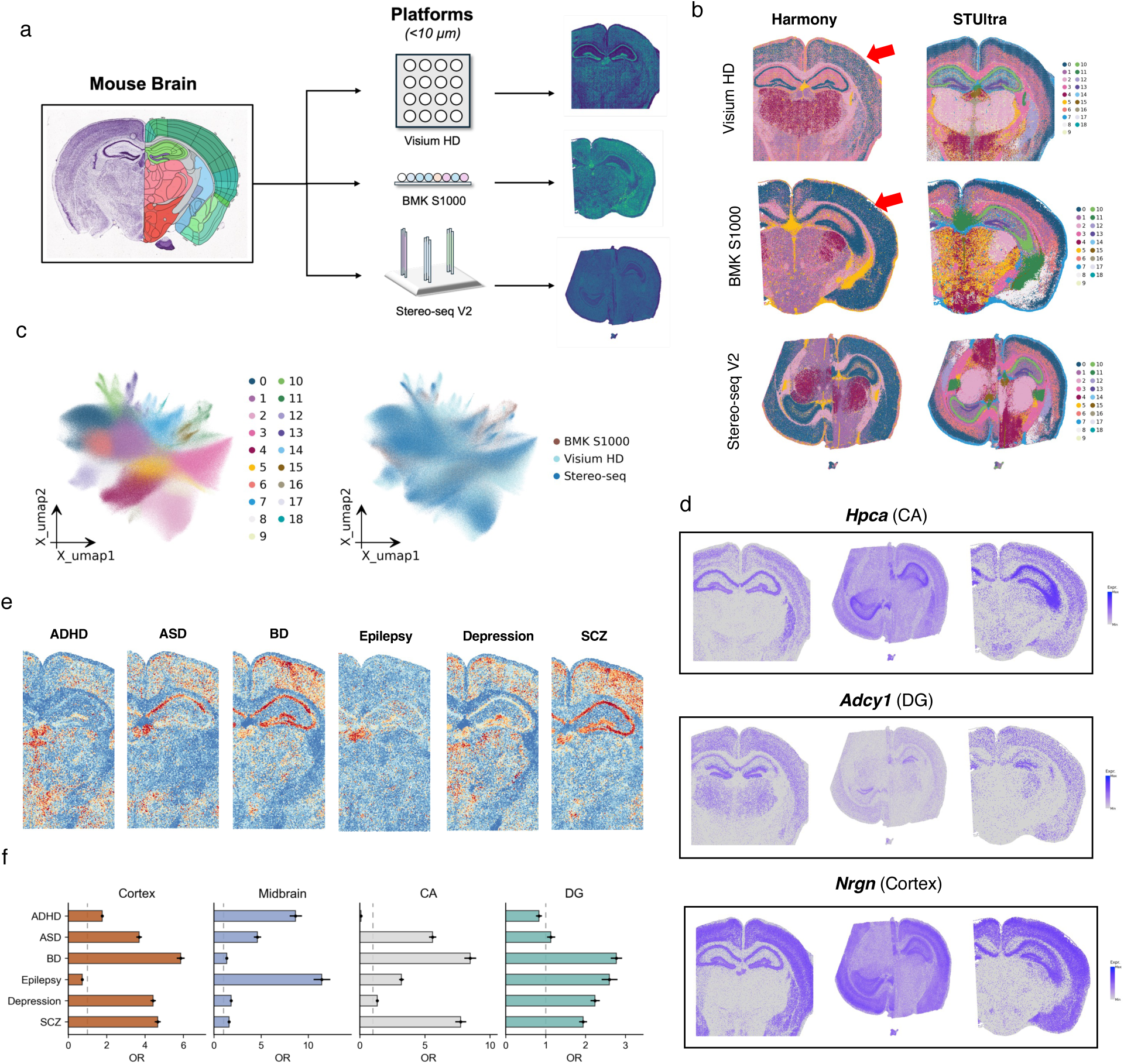
STUltra identified commonly shared domains from three different high-resolution platforms (Visium HD, BMK S1000, and Stereo-seq V2). (a) Three adult mouse brain datasets generated by three different platforms of ~ 10*μ*m spatial resolution: Visium HD, BMK S1000, and Stereo-seq V2. (b) Visualization results for spatial domains identified by integrating the three ST slices profiled via Harmony or STUltra. (c) UMAP plots of STUltra embeddings colored by slices (left) and by spatial domain (right). (d) Spatial expression patterns of selected markers for the cerebral cortex and hippocampus in the three ST slices of adult mouse brain. (e) Trait-mapping results for ADHD, Autism spectrum disorder (ASD), Bipolar disorder(BD), Epilepsy, depression and Schizophrenia(SCZ) with colours indicating the significance of association. (f) Odds ratios (ORs) of cortex, midbrain, CA and DG. Error bars represent the 95% confidence interval, with the centre indicating the OR. Bar plots ensure visibility of small-range error bars.

Many existing methods failed to complete correction on these ultra-large slices because of GPU memory limits or the high computational cost of their graph-based architecture constructions. Harmony completed the computation but could not effectively distinguish anatomically separate structures (Fig. 5b), so these anatomically separate regions were partially merged in the clusters generated by Harmony, leading to many spatial domains being poorly defined. In contrast, STUltra integrated spatial domains with clear boundaries that well align with the annotations from the Allen Mouse Brain Atlas, indicating effective recovery of the key structures across the platforms. From the three HD slices, we identified 19 spatial domains (as visualized by UMAP). Despite large differences in the unit size, the three datasets were perfectly aligned in the embedding space of the output data from STUltra (Fig. 5c). Notably, STUltra achieved consistent delineation for the major anatomical regions, despite large variations in the spatial resolution or in the transcript capture efficiency of the three platforms.

To validate the spatial domains identified by STUltra, we highlight the cerebral cortex and hippocampus in Fig. 5d as representative, atlas-grounded validations of cross-platform alignment. All integrated domains across the full sections are shown in Fig. 5b,c; As expected, the marker genes *Pltgds* and *Nrgn* were highly enriched in the cortical layers (Clusters 0 and 7), consistent with the known anatomical organization (Fig. 5d). Moreover, we observed distinct marker genes that were uniquely expressed in specific domains of the hippocampus, including *Hpca* in the CA region (Cluster 10) and *Adcy1* in the dentate gyrus (Cluster 13). Marker gene *Adcy1* showed a scoped and well-defined spatial pattern, further confirming the biological precision of STUltra’s integration results.

We also investigated whether the integrated cross-platform mouse brain landscape could bridge neuropsychiatric genetic associations to specific anatomical domains (Fig. 5e,f). When stratified by trait, the spatial enrichment analysis revealed distinct anatomical patterns of genetic association. ADHD and epilepsy showed the most focal profiles, with their strongest enrichment observed in the midbrain. Clinically, epilepsy is also frequently comorbid with ADHD, supporting overlapping neurodevelopmental mechanisms within deep brain structures [28]. In contrast, autism spectrum disorder (ASD), bipolar disorder (BD), and schizophrenia (SCZ) were most strongly enriched in hippocampal CA-associated clusters, indicating a shared convergence of psychiatric risk within specific hippocampal domains. Major depression showed a broader pattern, with prominent enrichment across cortical and dentate gyrus-associated clusters, consistent with a more distributed cortico-hippocampal contribution. Thus, although several neuropsychiatric traits shared enrichment within hippocampal and cortical territories, the dominant spatial signature differed by trait, highlighting trait-specific anatomical organization of genetic risk.

### 2.5. Long-term tracking of developmental domains in mouse embryo ST datasets

We conducted analysis on a Stereo-seq dataset of mouse embryos throughout eight developmental stages from E9.5 to E16.5, comprising of more than 500,000 cells in total (Fig. 4a). These data samples exhibit substantial size variation and strong temporal heterogeneity alongside the stages, ranging from small embryos at E9.5 with only a few thousand spatial bins to large E16.5 samples containing over 200,000 spatially resolved cells. After inter-stage integration, STUltra effectively mitigated batch effects arising from developmental differences, well-aligned spatial domains over time, and revealed coherent spatial correspondences throughout embryonic development. Importantly, the annotated tissues spanning the eight developmental stages displayed well-defined structural organization, accompanied by the expected expression patterns of known marker genes (Supplementary Fig. S10-S13).

At the individual tissues, we observed clear temporal correspondences that reflect continuous developmental dynamics (Fig. 4b). For instance, the heart remained morphologically stable throughout development, but gradually formed distinct chambers after E11.5; The liver was relatively small at E9.5 but expanded rapidly in later stages, with its hematopoietic function gradually shifting to the spleen by E15.5, as confirmed by the spatial enrichment of stage-specific marker genes (Supplementary Fig. S11). Skin development began at E12.5, when epidermal layers started to form and progressively enclose the embryo. STUltra also delineated unique-shaped structures, such as the meninges, which closed continuously and maintained consistent morphology during late embryogenesis.

STUltra also identified small yet functionally important structures such as the olfactory bulb, whose related gene expression emerged after E9.5, while the spatially coherent tissue structure became detectable only after E12.5. In addition, STUltra provided a continuous developmental perspective that differs from the original stage-specific annotations through long-range integration across embryonic stages (Fig. 4c). For example, the reference annotation assigns dermomyotome and muscle as distinct compartments, while STUltra grouped these regions into a unified cross-stage domain, reflecting their shared myogenic lineage and continuous developmental progression. The associated gene module reflects a skeletal muscle developmental program whose spatial expression tracks this pattern.

We further used trait-mapping results to evaluate the spatial distribution of genetic signals of the congenital heart disease (CHD) during mouse embryonic development (Fig. 4j-m). Based on the Louvain annotations of the final E16.5 slice, we selected six developmentally relevant modules— Cardiac, Mesenchymal/ECM, Neural crest/craniofacial, Circulatory, Epithelial/skin-associated, and Brain—and used the CHD spatial LDSC – log_10_(*P*) value as the strength of genetic association. Spot-level analysis on all the E16.5 modules showed that the Cardiac and Circulatory modules had the highest median CHD signals, with values of 0.734 and 0.730, respectively. Although the strongest CHD spatial signal appeared in the heart region, this association was not restricted to the heart; elevated signals were also detected in adjacent or developmentally related embryonic regions, including the Circulatory and Mesenchymal/ECM modules. This suggests that CHD genetic risk extends beyond the cardiac compartment and is associated with a broader embryonic developmental network involving cardiac neural crest migration, vascular and circulatory development, and early axial patterning programs. Along the E9.5–E16.5 developmental timeline, smoothed module-level signal curves revealed distinct spatiotemporal patterns of CHD-associated genetic signals, suggesting that disease risk may arise from coordinated developmental processes across multiple tissue structures rather than at a single cardiac lineage. Together, these results demonstrate that STUltra is a multi-function framework capable of not only reconstructing spatiotemporal development trajectories but also facilitating the understanding of stage-linked tissue organization and cross-stage correspondences.

### 2.6. Dissecting disease-associated spatial domains

To evaluate STUltra’s performance for disease analysis, we applied it to three related pathological settings involving disease-associated subtype identification, injury-to-repair spatial progression, and diseased-versus-healthy tissue remodeling, using Alzheimer’s disease, cardiac ischemic injury, and psoriasis as representative cases. Alzheimer’s disease (AD) is a progressive neurodegenerative disorder characterised by cognitive decline and widespread brain pathology. To investigate its underlying regulatory mechanisms, we applied STUltra to a mouse AD STARmap PLUS dataset containing both AD-model and age-matched control brains at 8 and 13 months of age [29]. Reference anatomical annotations of mouse brain regions in Fig. 6a provide an intuitive spatial framework for interpreting the results.

**Figure 6.**
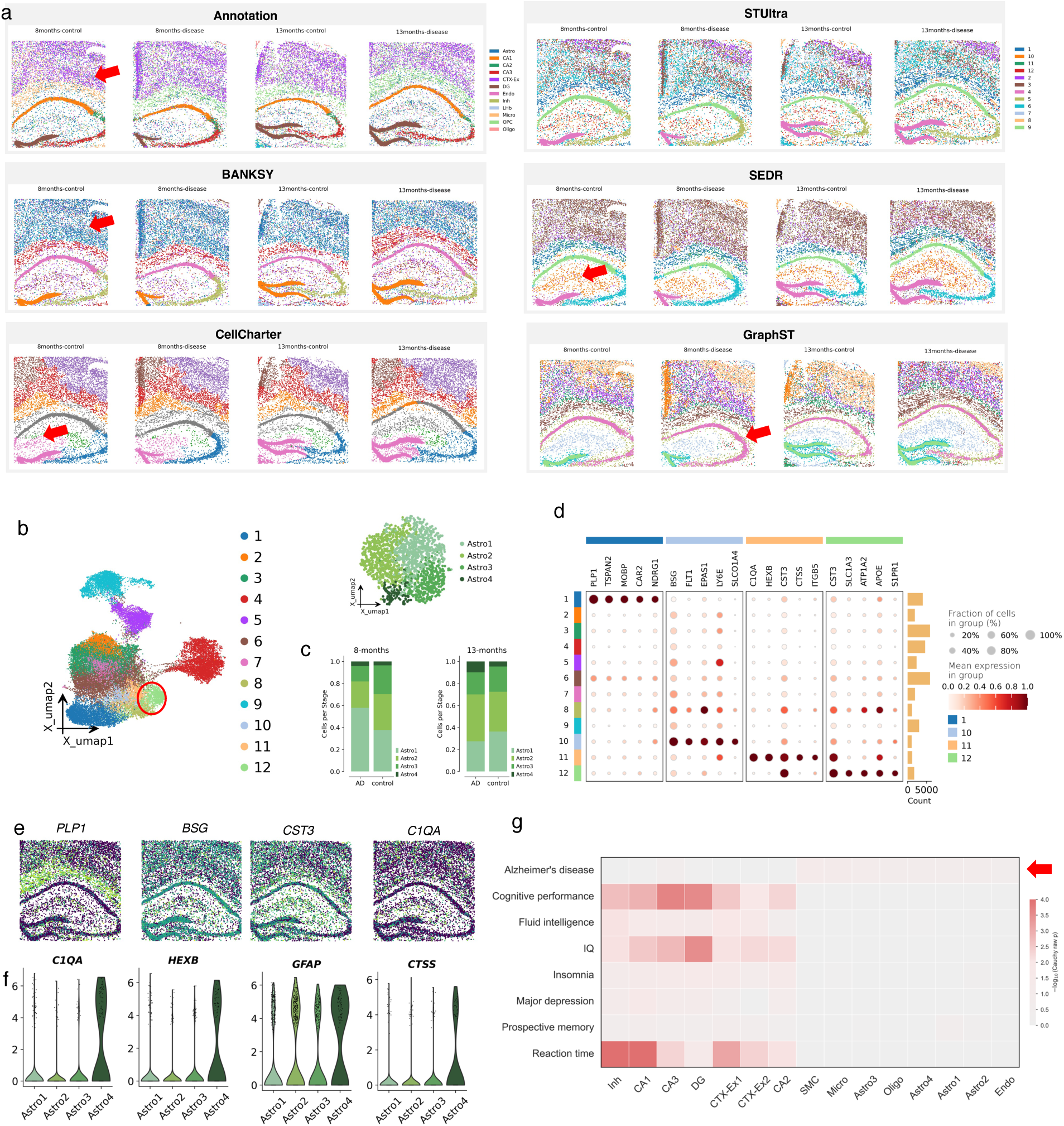
Analysis on the STARmap PLUS dataset, which contains omics data from brain slices of Alzheimer’s disease and control mice at 8 and 13 months of age. (a) Multi-method comparison of spatial domains identified by GraphST, CellCharter, BANKSY, SEDR, and STUltra on integrated brain sections, shown alongside the reference anatomical annotations. (b) UMAP visualization for all clusters, where cluster 12 (astrocytes) and its subtypes are highlighted. (c) Proportional changes of different astrocyte subtypes between AD and control samples at both ages. (d) Dot plot showing differential gene expression across twelve identified clusters. (e) Spatial expression maps of representative marker genes (PLP1, BSG, CST3, C1QA) identified by STUltra. (f) Violin plots showing the expression distributions of top marker genes in Astro4. (g) Tissue–trait association map. Columns represent tissue types, while rows represent traits. The color intensity indicates the statistical significance of the association between each tissue and trait.

After integrating these sections, we benchmarked STUltra against GraphST [9], CellCharter [18], SEDR [15], and BANKSY [16] (Fig. 6a). STUltra produced spatial domains with sharper tissue organisation and better preservation of local architecture for major brain cell populations, supporting finer dissection of disease-associated spatial patterns in AD. By contrast, GraphST, CellCharter, or BANKSY only partially separated AD and the control tissue but yielded coarser domain boundaries and missed fine-scale heterogeneity in spatial cellular organisation. These subtle differences are important for characterising microenvironmental remodelling during AD progression. The inferred STUltra clusters corresponded to known brain cell types, including astrocytes (Cluster 12), endothelial cells (Cluster 10), oligodendrocytes (Cluster 1), and microglia (Cluster 11). To understand more about astrocytic heterogeneity, we extracted all the astrocytes identified by STUltra and performed sub-clustering analysis, resulting in four distinct subpopulations: Astro1, Astro2, Astro3, and Astro4 (Fig. 6b-c).

The proportion of Astro4 increased significantly in the AD group in comparison with the controls. Although Astro4 only accounts for a very small fraction of the total cell population, they are indicators for dynamic astrocyte responses during disease progression (Fig. 6c). Compared with the other astrocyte subtypes, Astro4 showed marked up-regulation of C1QA, HEXB, GFAP and CTSS (Fig. 6f), consistent with the disease-associated astrocyte (DAA) phenotype described in the literature [30, 31]. Among these genes, C1QA showed the most prominent up-regulation in subtype Astro4 (Fig. 6e,f). This gene has been linked to aberrant microglial activation and synaptic degeneration that contribute to cognitive decline in AD [32, 33].

We compared AD with cognitive and neuropsychiatric traits to determine whether its genetic association pattern occupied distinct spatial cell-state contexts (Fig. 6g). First, the AD mapping was biased toward non-neuronal cell types, with the strongest emphasis on Micro and Astro4, while cognitive-related traits, including IQ, cognitive performance, fluid intelligence, and reaction time, were mainly mapped onto neuronal and hippocampal regions such as CA1, CA3, and DG. On the other hand, the AD Cauchy signals were not prominent in these neuronal regions but were relatively stronger in Micro and Astro4. This observation suggests that AD genetic risk in this spatial dataset is enriched in microglia-mediated immune responses and Astro4-related reactive astrocyte states, distinguishing it from cognitive traits dominated by neuronal synaptic or hippocampal circuit signatures. Meanwhile, the relative proximity between prospective memory and AD (cosine = 0.356) suggests that part of cognitive decline may be linked to AD-related pathological processes.

Beyond identifying static disease-associated cell states, STUltra also captured dynamic spatial reorganization in a mouse cardiac ischemic injury dataset, resolving border, infarct, and remote zones and ordering injury-responsive molecular transitions spanning post-injury time points (Supplementary Fig. S3, Fig. S16). In pair-matched psoriatic lesional and non-lesional skin, STUltra identified lesion-expanded spatial domains with psoriasis-associated marker expression and immune-related pathway enrichment (Supplementary Fig. S4). Together, these three disease case studies demonstrate the versatility of STUltra in dissecting spatial patterns across diverse pathological contexts, including chronic neurodegeneration, acute tissue injury, and autoimmune inflammation, each revealing finer-grained spatiotemporal heterogeneity.

## 3. Discussion

We developed STUltra, a scalable framework that integrates multi-sample ST data and links tissue domain organizations to GWAS-derived trait associations. By leveraging interval-sampled integrative hypergraph construction together with a robust graph autoencoder and contrastive learning, STUltra can organize intra-slice spatial neighborhoods and inter-slice biological correspondences effectively into a structure-boundary-distinct landscape. This design has improved performance in preserving true biological variation, enabled spatiotemporal trait-cell mapping across heterogeneous tissue contexts, and reduced dependence on exhaustive graph construction used by graph-based methods such as STAligner and SEDR.

Regarding downstream applications, STUltra highlights its utility as a versatile tool for advancing ST data analysis and interpretation. In the sagittal mouse brain sections, the reconstruction of the hippocampal trisynaptic circuit and cortical laminar patterns provided the spatial context in which epilepsy-associated signals were more focally localized to forebrain, hippocampal, and subcortical seizure-related structures. In the cross-platform adult brain integration, the preservation of hippocampal and cortical domains enabled schizophrenia-associated signals to point to hippocampal CA-associated domains despite differences in spatial resolution and capture efficiency. From the mouse embryo datasets spanning eight developmental time points, STUltra aligned fragmented domain labels and revealed continuous tissue dynamics while showing that congenital heart disease signals extended beyond cardiac regions to circulatory and mesenchymal/ECM developmental modules. About the colorectal cancer study, the alignment of tumor-adjacent local spatial structures placed Crohn’s disease signals in myeloid-enriched microenvironments, newly linking genetic risk to immune-inflamed tissue niches. In the Alzheimer’s disease sections, the identification of reactive astrocyte subtypes further showed that genetic association was biased toward Micro/Astro4 reactive glial contexts rather than neuronal hippocampal regions. Overall, STUltra provides a unified framework for resolving spatially organized cell populations and mapping trait-associated tissue structures across organ development, disease progression, and tissue maintenance.

There are some limitations of STUltra and areas for future improvement and study. First, integrating histological or high-resolution imaging data could further enhance the spatial alignment and improve the biological interpretability of the identified spatiotemporal domains. Second, the current framework primarily focuses on transcriptomic integration; extending it to incorporate additional modalities, such as proteomics or metabolomics data, may provide a more comprehensive view of spatial regulation. In addition, GWAS-based trait mapping should be interpreted as association-level prioritization of candidate spatial niches, because it depends on GWAS quality, SNP-gene mapping, and domain-level aggregation, and does not by itself establish causality. In the future, adding more steps of analysis within the STUltra framework may further enhance its ability to characterize complex biological processes and disease progression, such as RNA velocity inference and cell–cell interaction modeling.

## 4. Methods

### 4.1. Preprocessing

STUltra uses gene expression data and spatial coordinates from multiple ST slices as input. Throughout the Methods, a *spatial input unit* denotes one row of the input expression matrix with associated spatial coordinates. Depending on the dataset, this unit can be a Visium capture spot, high-resolution bin or bead, segmented cell, or another aggregated spatial expression unit; STUltra does not require the unit to be an individual transcript molecule. For each slice, the raw gene expression counts are normalized and log-transformed using the `Scanpy` package [6]. Subsequently, the top 3,000–5,000 highly variable genes (HVGs) are selected depending on the integration setting. To ensure consistency across the slices, we use the intersection of HVGs as the shared feature set for downstream analysis.

### 4.2. Main Steps Taken by STUltra

STUltra consists of three main components: (i) interval-sampled (interval-gridded) integrative hypergraph construction for capturing global-local structures from ultra-large multi-slice data; (ii) robust graph autoencoder (GAE) for representation learning from its sparse graph representation; (iii) contrastive learning for batch integration (Supplementary Alg. 1). The training process has two stages. In the first stage, the robust GAE is pre-trained to learn spatially consistent embeddings from individual slices; in the second stage, the model is fine-tuned with the contrastive objective to align across the slices to achieve integrated embeddings.

#### Interval-sampled integrative hypergraph construction

By interval sampling, STUltra constructs an integrative hypergraph to jointly describe two types of neighborhood relations that are required for multi-slice integration. The first type is the intra-slice neighborhood hyperedge, which links a sampled spatial input unit with its local spatial context within the same tissue section. The second type is the inter-slice neighborhood hyperedge, which links biologically corresponding local contexts across different slices and is subsequently refined by the contrastive alignment procedure. Together, these intra- and inter-slice hyperedges provide a unified neighborhood construction for preserving tissue topology and aligning related domains across slices.

#### Interval sampling

To scale up the integration scalability of our method for arbitrarily large ST slices, we take an interval sampling strategy with a fixed spatial stride. Sampling is carried out by scanning the tissue slice with a regular grid to select representative spatial input units through nearest-neighbor matching. Given a tissue slice consisting of *N* spatial input units represented as

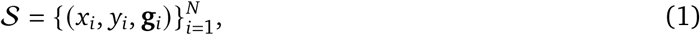

we use a fixed spatial interval length Δ *>* 0 to construct a regular grid aligned with the slice boundary. Specifically, we compute the lower bounds *x*_min_ = min_*i*_ *x*_*i*_ and *y*_min_ = min_*i*_ *y*_*i*_, and create grid points

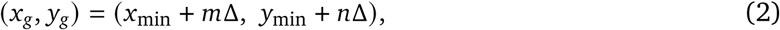

where, *m*, *n* are non-negative integers such that (*x*_*g*_, *y*_*g*_) lies within the spatial bounding box of the slice. For each grid point, we query the nearest observed spatial input unit in its Euclidean space and select the corresponding gene expression vector. When multiple grid points are mapped to the same spatial input unit, duplicate selections are removed. This results in a spatially uniform subset of spatial input units (called a *patch*) that approximates regular sampling over the tissue area. The number of sampled spatial input units scales approximately with the tissue area divided by Δ^2^, enabling efficiency in the subsequent processing of the ultra-large ST slices. Our interval mapping strategy typically samples 4 or 9 times on the same slice, creating a different subset of spatial input units (or a different patch) each time. These interval-sampled patches provide the scalable units for the subsequent integrative hypergraph construction, in which local neighborhoods within slices and corresponding neighborhoods across slices are organized in a unified manner.

If the omics data from the slice is considered as a 2-dimensional image matrix, then our interval mapping can be conceptually interpreted as a dilated convolutional operator [34] on a regular image grid. Here, the fixed spatial interval corresponds to the dilation rate, and the operation resembles applying a Δ × Δ convolutional kernel whose elements are all zero except for the center element being 1. The multiple convolution-transformed patches of the original ultra-large slice are then integrated by *batch training* to signify embeddings from the slice. This is exactly the one-facet meaning of STUltra’s ‘multi-integration’; and another-facet meaning of ‘multi-integration’ is to process omics data from multiple tissue slices. So, it is a double-looped integration.

#### Why interval-based batch training is effective

From an optimization perspective, interval mapping enables batch-based training to approximate full-slice learning in a controlled manner. By uniformly scanning the spatial domain with a fixed stride, the sampled subset provides a near-uniform coverage of the tissue area to ensure that local spatial patterns and global structural statistics are preserved in expectation. When model updates are computed at such sampled subsets, the resulting gradients can be viewed as low-variance approximations of the gradients obtained from the full slice [35, 36]. In the ST setting, interval mapping further reduces redundancy among neighboring spatial input units, which improves sample efficiency without introducing strong spatial bias. As training proceeds, the sampled representation increasingly reflects the structure of the full multi-slice neighborhood system, leading to optimization dynamics that are similar to those observed under full-slice training. Empirically and theoretically, this type of structured sampling has been shown to preserve model performance while significantly reducing computational cost, as also observed in large-scale graph learning theories [37].

#### Projection to sparse graph representation

For efficient graph autoencoding, the high-order neighborhood relations from the integrative hypergraph are represented as sparse pairwise adjacency. Within each slice, neighbors are determined either by Euclidean distance or by the *k*-nearest neighbors (kNN), where the parameter *k* or radius *r* can be adjusted to adapt to different ST scenarios and platforms. If nodes *i* and *j* participate in the same projected intra-slice neighborhood relation, we set *A*_*ij*_ = *A*_*ji*_ = 1. The adjacency matrix *A* is independently constructed for each sampled slice representation and then concatenated into a unified graph matrix, stored efficiently in sparse format.

#### GPU-accelerated graph preprocessin

To further improve the efficiency of the projected neighborhood construction and neighbor querying, we add several `torch_geometric` components into the preprocessing pipeline. We use `torch_cluster.knn_graph` to build kNN edges in the GPU using the spatial coordinates of each slice, which directly produces PyG-compatible `edge_index`. The gene expression features and coordinates are then packed into `torch_geometric.data.Data` objects, and multiple slices are combined using `torch_geometric.data.Batch.from_data_list` for downstream training. To reduce memory and copying overhead, we convert the adjacency structure into a `torch_sparse.SparseTensor` in CSR or CSC form and keep it in the GPU, which enables constant-time neighbor access and efficient sparse matrix operations. For aggregation functions such as neighbor-wise summation or maximization, we rely on GPU implementations to achieve high throughput and noticeable speedup.

#### Robust graph autoencode

The latent representation learning at spatial gene graph is achieved through a robust graph autoencoder, which consists of an autoencoder, a node discrimination module, and an uncertainty module (Fig. 1).

#### Encoder

The encoder is constructed with *L* graph attention (GAT) layers and takes the spatial graph *G* = (*X*, *E*) as input, where *X* denotes the normalized gene expressions and *E* denotes the corresponding graph adjacency structure. For the *k*-th encoder layer (*k* ∈ {1, 2*,. . ., L* – 1}), the output representations of the nodes are computed using an attention mechanism, formulated as

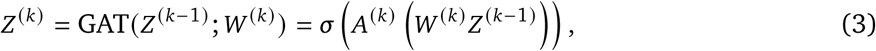

where *W*^(*k*)^ is the trainable weight matrix, *σ* is the nonlinear activation function, and *Z*^(0)^ = *X* represents the initial node features of the spatial graph *G*. The attention matrix *A*^(*k*)^ is obtained following the graph attention approach [38], which computes pairwise attention coefficients to adaptively aggregate neighborhood information based on both feature similarity and spatial adjacency. The latent embedding *Z*^(*L*)^ is finally calculated by the *L*-th layer in the encoder with a simple projection

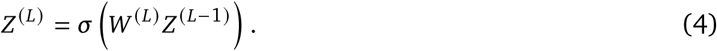

#### Decoder

To reconstruct the expression features, the decoder is designed to have the same number of *L* layers as the encoder, following a symmetric but reversed propagation process. The decoding process is defined as:

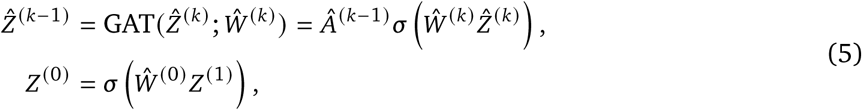

where *k* ∈ {1, 2*,. . ., L* – 1}, *Ŵ* ^(*k*)^ denotes the trainable weight matrix in the *k*-th decoder layer, and *Â*^(*k*–1)^ represents the attention matrix.

To prevent overfitting and maintain consistency in the autoencoder, we typically set *Ŵ* ^(*k*)^ = *W*^(*k*)^ and *Â*^(*k*)^ = *A*^(*k*)^ for all the layers. The decoder progressively restores the spatially smoothed gene expression features from the latent embeddings, yielding *Ẑ*^(0)^ as the final reconstructed expression matrix. Through stacked GAT layers, the reconstruction process is performed by minimizing the mean squared error loss:

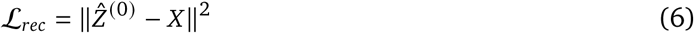

#### Probabilistic uncertainty estimation

ST data from different biological conditions often exhibit uncertain batch effects, making it difficult for the model to generalize across heterogeneous tissue domains. To enhance robustness, we introduce an Uncertainty Estimation (UE) module into the GAE framework. Instead of treating each feature statistic as a deterministic value, UE assumes that the mean and standard deviation should follow probabilistic distributions reflecting potential spatial or technical variations [39]. Specifically, for each feature channel, the mean and standard deviation are modeled as Gaussian random variables 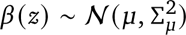 and 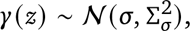 respectively. The uncertainty scales ε_*μ*_ and ε_*σ*_ are estimated at the mini-batch level as

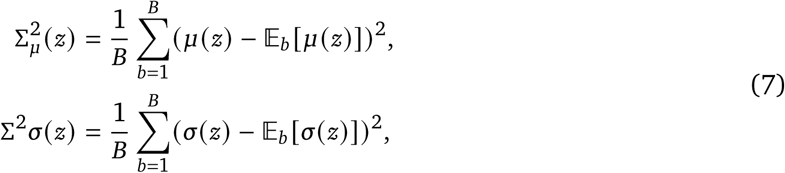

where *B* is the batch size. Random perturbations are then introduced via reparameterization

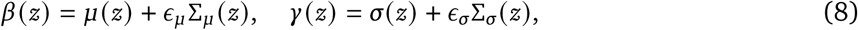

with *ϵ*_*μ*_, *ϵ*_*σ*_ ~ *N* (**0**, **1**).

The adjusted representation can be obtained through

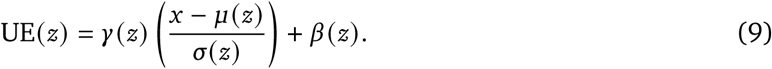

This module is flexibly incorporated after each layer of the GAE encoder and decoder during training to simulate potential batch effects among heterogeneous ST slices. To better illustrate this process, we also provide a diagram visualization in Supplementary Fig. S5.

#### Node discrimination module

For ultra-large ST slices, capturing local spatial neighborhoods is a crucial step because neighboring cells often share strong molecular correlations that may be lost when only global representations are considered. To enhance such potential of capturing local patterns in the GAE, we apply a node discrimination module following Liu et al. [40]. Given *G* = (*X*, *E*), we generate an augmented view *G*^′^ by applying graph augmentations *τ* (e.g., attribute noise, edge dropout). The shared encoder with projection head *f* maps *G* and *G*′ into node embeddings *Z* and *Z*′, respectively.

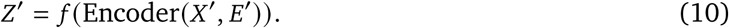

Then, the binary cross-entropy loss is applied to separate the original nodes from the augmented ones

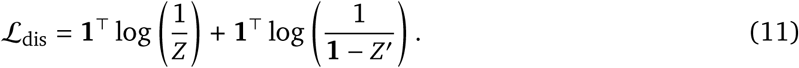

where **1** ∈ ℝ^*N*×1^.

In the first stage, the autoencoder is trained to preserve both global and local structures by minimizing the following loss

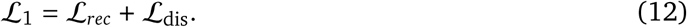

#### Contrastive learning

Integrating multiple ultra-large ST slices presents tremendous challenges for the embedding alignment across the slices, especially when each contains up to one million spatial input units. Our interval-sampled integrative hypergraph construction can reduce the total number of spatial input units processed per training step while preserving both intra-slice local neighborhoods and inter-slice correspondence neighborhoods, providing manageable subsets for downstream embedding and contrastive learning.

The sparse graph representations projected from the integrative hypergraph capture global-local relationships, which are then encoded into latent embeddings used in the contrastive learning stage. To well align embeddings spanning such slices while mitigating batch effects, we adopt a contrastive learning strategy based on the InfoNCE loss [41]. InfoNCE maximizes similarity for positive pairs while contrasting them against a set of negatives.

Here, we extend contrastive learning to the cross-slice ST setting by first constructing inter-slice neighborhood hyperedges from embedding-space similarity, where explicit one-to-one correspondences between spatial input units are unavailable. Given an anchor spatial input unit with index *i* in slice *s*, we define its positive counterpart index *j* as that of the most similar spatial input unit identified from the other slices via mutual nearest neighbors (MNNs), while negatives are randomly sampled from the same slice as the anchor. In this sense, MNN-derived positive pairs serve as projected inter-slice neighborhood relations that connect biologically corresponding local contexts across slices. The InfoNCE loss in batch-disentangled contrastive learning is then formulated as

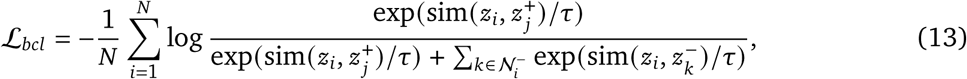

where sim(·) denotes the cosine similarity, *z*_*i*_ and *z*_*j*_^+^ are the latent embeddings of the anchor and its positive counterpart, *N*_*i*_^-^ represents the set of negative samples from the same slice as *i*, and *τ* is a temperature parameter. The final objective in the second stage is given as

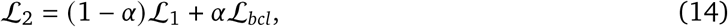

where *α* is a hyperparameter that controls the degree of multi-slice integration and is set to 0.2 by default. Moreover, this contrastive loss enables efficient optimization using batch training techniques on ultra-large ST slices.

For ultra-large ST slices, each of them contains dense local gene expression profiles, which may dominate the alignment process if not properly controlled. This batch-disentangled design ensures that anchors are aligned with positives to achieve cross-slice integration, while negatives prevent the model from overfitting to dense local structures, allowing the network to integrate heterogeneous slices effectively without being dominated by intra-slice patterns.

#### The overall process

We train the model through an iterative optimization process that alternates between embedding learning and cross-slice pairing refinement under the batch-disentangled contrastive framework. In the first stage, the robust GAE is pre-trained to obtain initial embeddings. Then, the encoder is fine-tuned using the batch-disentangled pairs jointly with contrastive learning, thereby improving both alignment and representation consistency. After each update, new contrastive pairs are recomputed based on the refined embeddings, and the process is repeated until convergence or a fixed number of iterations is reached. The final embeddings provide a unified representation for downstream spatial integration and analysis. Detailed pseudocode can be found in Alg. 1.

### 4.3. Differential expression and GO enrichment analysis

DEGs for the spatial domains in the condition integration were identified using the Wilcoxon test implemented in the `rank_genes_groups()` function of the Python package `Scanpy` (v1.9.3), with an adjusted *P* value (Benjamini–Hochberg correction) threshold of 0.01. GO enrichment analysis of the domain-specific DEGs was conducted using the `bulk.geneset_enrichment` module of the Python package `Omicsverse` (v1.7.7).

### 4.4. Benchmark methods for ST data integration

To evaluate the integration performance, we compared ours with the following methods: Harmony [5], STAligner [17], STitch3D [14], SEDR [15], SpaBatch [20], SpaGIC [42], BANKSY [16], and CellCharter [18]. In addition, we used Scanpy [6] to generate uncorrected embeddings from the combined raw input data (Tab. 1).

**Table 1.** Comparison of benchmarked integration methods.

| Method | Key Features | Reference | Year |
| --- | --- | --- | --- |
| Scanpy | Uncorrected raw embeddings | Ref. [6] | 2018 |
| Harmony | Multi-sample, PCA-based, soft clustering | Ref. [5] | 2019 |
| STAligner | Graph attention, triplet loss | Ref. [17] | 2023 |
| STitch3D | 3D reconstruction, global graph | Ref. [14] | 2023 |
| CellCharter | Graph-based clustering, neighborhood-aware | Ref. [18] | 2024 |
| SEDR | GNN, PCA features, Harmony | Ref. [15] | 2024 |
| BANKSY | Spatial smoothing, neighborhood aggregation | Ref. [16] | 2024 |
| SpaGIC | Graph-informed clustering, contrastive learning | Ref. [42] | 2024 |
| SpaBatch | Masked autoencoders, deep embedded clustering | Ref. [20] | 2025 |

#### Harmony

Harmony is an integration method designed for multi-sample or multi-batch single-cell and spatial omics data. It projects the data with PCA and iteratively adjusts embeddings to reduce batch effects while preserving biological variation. It uses soft clustering with a diversity penalty to ensure that clusters from multiple sources are achieved, thereby achieving cross-batch alignment.

#### STAligner

STAligner is a method for aligning and integrating multiple spatial transcriptomics datasets. It leverages graph attention neural networks to encode spatial correlation and employs a triplet loss to achieve integration. We tested STAligner under its default parameter settings.

#### STitch3D

STitch3D integrates multiple 2D slices into a 3D reconstruction. It first aligns slices in space and then constructs a global 3D graph that incorporates inter-slice spatial relations. A GNN is then applied to learn latent representations, achieving both batch correction and spatial integration.

#### SEDR

SEDR is a spatial embedding representation method based on GNN, with PCA embeddings used as node features. We followed the original settings provided in the official tutorial for unsupervised batch correction experiments, where the latent embeddings are further processed by Harmony.

#### SpaGIC

SpaGIC is a graph-based deep learning framework that leverages spatial neighborhood details via graph and contrastive learning. It outperforms existing methods in key spatial transcriptomics tasks, including multi-slice analysis.

#### SpaBatch

SpaBatch is a multi-slice integrative framework using masking, two-stage training (VGAE pretraining + DEC fine-tuning), and triplet contrastive learning to handle noise, batch effects, and inter-slice variability. It aligns cross-slice biological structures while preserving discriminative features, enabling accurate 3D spatial domain identification.

#### BANKSY

BANKSY augments each cell’s transcriptome with spatial neighborhood summaries, jointly encoding cell state and microenvironment. BANKSY enables unified cell typing and spatial domain segmentation across platforms. For integration, BANKSY embeddings may be batch-corrected by Harmony.

#### CellCharter

CellCharter is a Python tool for spatial cell niche identification and comparison across samples. It learns batch-corrected embeddings with scVI/scArches and assigns cells to domains using neighborhood features and Gaussian mixture models. CellCharter supports domain stability and niche enrichment analyses.

### 4.5. Implementation details of STUltra

The STUltra encoder is composed of two GAT layers with hidden dimensions 512 and 30, while the decoder mirrors the encoder structure. Model optimization is performed using the AdamW optimizer [43] with a learning rate of 10^-4^ and weight decay of 10^-4^. The robust graph autoencoder and the search process of MNN pairs are implemented using the PyTorch Geometric library [44]. The training proceeds in two phases: first, the autoencoder is trained to produce initial embeddings; next, embedding refinement and batch-disentangled contrastive training are alternated every 100 iterations until the total number of iterations is reached. All hyperparameters were optimized based on their performance on the DLPFC dataset. The default number of iterations is 500 (or 200) for the HD platforms.

### 4.6. Clustering and evaluation

The spatial domains were identified by clustering the embeddings obtained from STUltra. For datasets with prior knowledge of the number of domain labels, we applied the `mclust` clustering algorithm [45] implemented in the R package `mclust` (v5.4.9). For datasets without such prior knowledge, we used the Louvain algorithm [46] implemented in the Python package `Scanpy` (v1.9.3).

To evaluate clustering accuracy, we adopted the Adjusted Rand Index (ARI) [21], which measures the similarity between the clustering result and the reference annotation. Let *RI* denote the Rand Index, and *E*[*RI*] its expected value under random labeling. Then ARI is given as

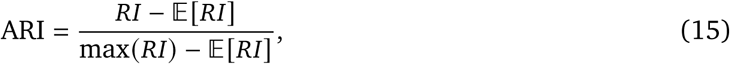

where *RI* is Rand index. ARI takes a maximum value of 1 when the two partitions are identical, and has an expected value of 0 for random cluster assignments.

We also used the Normalized Mutual Information (NMI) [47] to assess clustering performance, which quantifies the mutual information between the clustering result and reference annotation, normalized by the average entropy of the two partitions. Let *H*(*X*) and *H*(*Y*) represent the entropies of the reference annotation and clustering result, respectively, and *I* (*X*, *Y*) denote their mutual information. Then, NMI is defined as

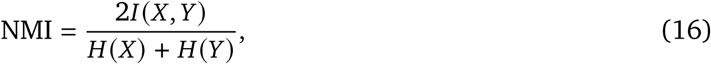

where 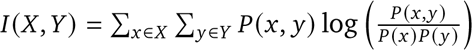 is the mutual information, with *P*(*x*, *y*) being the joint probability distribution of *X* and *Y*, and *P*(*x*), *P*(*y*) their marginal distributions. NMI ranges from 0 (no mutual information) to 1 (perfect agreement between partitions).

### 4.7. Traits mapping onto spatiotemporal domain landscapes

To link ST integration with human complex traits, we coupled the STUltra-integrated spatial representations with GWAS summary statistics following the main strategy of gsMap [22]. The same integrated spatial input units and STUltra-derived or annotated domains used for tissue interpretation were used as trait-mapping units, allowing trait signals to be compared across slices, sequencing platforms, developmental stages, or disease conditions within a shared representation space.

For each integrated spatial domain, local expression patterns after STUltra integration were summarized into gene-level spatial scores, which quantify the relative spatial specificity of each gene in the corresponding spatial context. These gene-level scores were assigned to SNP annotations through SNP-gene mapping, and stratified linkage disequilibrium score regression (S-LDSC) was used to test whether SNPs with higher spatial scores explained disproportionate trait heritability. For spatial domains or annotated regions, point-wise association *P* values were aggregated using the Cauchy combination test, yielding a domain- or region-level significance score for each trait. The resulting scores were compared across tissue structures to prioritize trait-associated spatial niches and to interpret how genetic associations were distributed over the integrated tissue landscape. Because this procedure is based on GWAS summary statistics and spatially aggregated expression profiles, the resulting maps represent association-level evidence rather than causal validation.

### 4.8. Datasets description

We collected a diverse pool of spatial transcriptomic datasets originating from multiple tissues, platforms, or species to evaluate the performance of our method. The human DLPFC dataset profiled by a 10x Visium platform consists of four adjacent tissue slices (151507-151510), containing ~ 4, 000 input units. The mouse brain dataset includes two sagittal slices profiled by 10x Visium, having 2,695 (anterior) or 3,355 (posterior) input units. More importantly, we analyzed three high-resolution mouse brain datasets: a 10x Visium HD slice (579,500 input units), a BMK S1000 slice (249,130 input units), and a Stereo-seq v2 slice (577,094 input units) [19]. The mouse embryo dataset profiled by Stereo-seq covers developmental stages from E9.5 to E16.5, with each stage creating 5, 000 ~ 120, 000 input units. The human colorectal cancer dataset was profiled by the 10x Visium HD platform, consisting of two slices (P1 and P2) each having 500,000 or more input units. The human skin dataset profiled by 10x Visium contains two slices (Non-lesional and Lesional) containing 955 or 654 input units [48]. The mouse heart dataset profiled by 10x Visium includes four slices corresponding to different myocardial infarction (MI) time points: 1 hour, 4 hours, day 3, and day 7 [49]. Each slice contains an input unit count ranging from 2,236 to 2,578. A detailed summary of all these datasets is provided in Supplementary Tab. S1.

### 4.9. Computing Environment

STUltra is implemented using `Pytorch` (v2.4.0) and `Python` (v3.9). All the experiments were performed on a server with an Intel Xeon(R) Platinum 8360Y CPU and an NVIDIA GeForce RTX 4090 GPU (24 GB memory).

## 5. Data availability

Source data for all the figures are available. The datasets analyzed by this study are all from publicly available websites (Supplementary Table S1). Specifically, the human DLPFC dataset can be accessed in the spatialLIBD package (http://spatial.libd.org/spatialLIBD). The mouse sagittal posterior and anterior brain data can be accessed at https://support.10xgenomics.com/spatial-gene-expression/datasets/1.0.0/V1_Mouse_Brain_Sagittal_Posterior and https://support.10xgenomics.com/spatial-gene-expression/datasets/1.0.0/V1_Mouse_Brain_Sagittal_Anterior, respectively. The mouse brain data generated by Visium HD, BMK S1000, and Stereo-Seq v2 are available at the website (www.genographix.com). The mouse embryo data can be accessed at https://db.cngb.org/stomics/mosta/. The human colorectal cancer Visium HD data are available from 10X Genomics data website: https://www.10xgenomics.com/datasets/visium-hd-cytassist-gene-expression-libraries-of-human-crc. The human lesional and nonlesional skin data are publicly available at GEO under the accession number GSE202011. The mouse heart with myocardial infarction and mechanical injury data can be downloaded from GEO database with accession code GSE214611, and the metadata can be downloaded from https://drive.google.com/file/d/161iiznh3I8eiLe9tqb2xU7QDFmo6hLWD/view?usp=sharing.

## 6. Code availability

An open-source Python implementation of the STUltra package is available at https://github.com/ZZhangsm/STUltra.

## Acknowledgements

This study was supported by grants from the NSFC Project W2531048 and the 111 Project D23017.

## References

[1] Kristen R Maynard, Leonardo Collado-Torres, Lukas M Weber, Cedric Uytingco, Brianna K Barry, Stephen R Williams, Joseph L Catallini, Matthew N Tran, Zachary Besich, Madhavi Tippani, et al. Transcriptome-scale spatial gene expression in the human dorsolateral prefrontal cortex. Nature neuroscience, 24(3):425–436, 2021.

[2] 10x Genomics. Xenium in situ. https://www.10xgenomics.com/platforms/xenium, 2024. Accessed: 2024.

[3] Michelli Faria de Oliveira, Juan Pablo Romero, Meii Chung, Stephen R Williams, Andrew D Gottscho, Anushka Gupta, Susan E Pilipauskas, Seayar Mohabbat, Nandhini Raman, David J Sukovich, et al. High-definition spatial transcriptomic profiling of immune cell populations in colorectal cancer. Nature Genetics, pages 1–12, 2025.

[4] Ao Chen, Sha Liao, Mengnan Cheng, Kailong Ma, Liang Wu, Yiwei Lai, Xiaojie Qiu, Jin Yang, Jiangshan Xu, Shijie Hao, et al. Spatiotemporal transcriptomic atlas of mouse organogenesis using dna nanoball-patterned arrays. Cell, 185(10):1777–1792, 2022.

[5] Ilya Korsunsky, Nghia Millard, Jean Fan, Kamil Slowikowski, Fan Zhang, Kevin Wei, Yuriy Baglaenko, Michael Brenner, Po-ru Loh, and Soumya Raychaudhuri. Fast, sensitive and accurate integration of single-cell data with Harmony. Nature methods, 16(12):1289–1296, 2019.

[6] F Alexander Wolf, Philipp Angerer, and Fabian J Theis. SCANPY: large-scale single-cell gene expression data analysis. Genome biology, 19(1):15, 2018.

[7] Joyce B Kang, Aparna Nathan, Kathryn Weinand, Fan Zhang, Nghia Millard, Laurie Rumker, D Branch Moody, Ilya Korsunsky, and Soumya Raychaudhuri. Efficient and precise single-cell reference atlas mapping with symphony. Nature communications, 12(1):5890, 2021.

[8] Anjali Rao, Dalia Barkley, Gustavo S França, and Itai Yanai. Exploring tissue architecture using spatial transcriptomics. Nature, 596(7871):211–220, 2021.

[9] Yahui Long, Kok Siong Ang, Mengwei Li, Kian Long Kelvin Chong, Raman Sethi, Chengwei Zhong, Hang Xu, Zhiwei Ong, Karishma Sachaphibulkij, Ao Chen, et al. Spatially informed clustering, integration, and deconvolution of spatial transcriptomics with graphst. Nature Communications, 14(1):1155, 2023.

[10] Tian Tian, Jie Zhang, Xiang Lin, Zhi Wei, and Hakon Hakonarson. Dependency-aware deep generative models for multitasking analysis of spatial omics data. Nature Methods, 21(8): 1501–1513, 2024.

[11] Yuqiao Gong, Xin Yuan, Qiong Jiao, and Zhangsheng Yu. Unveiling fine-scale spatial structures and amplifying gene expression signals in ultra-large st slices with HERGAST. Nature Communications, 16(1):1–14, 2025.

[12] Ron Zeira, Max Land, Alexander Strzalkowski, and Benjamin J Raphael. Alignment and integration of spatial transcriptomics data. Nature Methods, 19(5):567–575, 2022.

[13] Bingjie Dai, Litai Yi, Peizhuo Wang, Hanshuang Li, Pengwei Hu, Yancheng Song, Jixiang Xing, Zhenxing Feng, Zhiyuan Yuan, and Yongchun Zuo. 3d-ot: a deep geometry-aware framework for heterogeneous slices alignment of spatial multi-omics. Nature Methods, pages 1–12, 2026.

[14] Gefei Wang, Jia Zhao, Yan Yan, Yang Wang, Angela Ruohao Wu, and Can Yang. Construction of a 3D whole organism spatial atlas by joint modelling of multiple slices with deep neural networks. Nature Machine Intelligence, 5(11):1200–1213, 2023.

[15] Hang Xu, Huazhu Fu, Yahui Long, Kok Siong Ang, Raman Sethi, Kelvin Chong, Mengwei Li, Rom Uddamvathanak, Hong Kai Lee, Jingjing Ling, et al. Unsupervised spatially embedded deep representation of spatial transcriptomics. Genome Medicine, 16(1):12, 2024.

[16] Vipul Singhal, Nigel Chou, Joseph Lee, Yifei Yue, Jinyue Liu, Wan Kee Chock, Li Lin, Yun-Ching Chang, Erica Mei Ling Teo, Jonathan Aow, et al. Banksy unifies cell typing and tissue domain segmentation for scalable spatial omics data analysis. Nature genetics, 56(3):431–441, 2024.

[17] Xiang Zhou, Kangning Dong, and Shihua Zhang. Integrating spatial transcriptomics data across different conditions, technologies and developmental stages. Nature Computational Science, 3 (10):894–906, 2023.

[18] Marco Varrone, Daniele Tavernari, Albert Santamaria-Martínez, Logan A Walsh, and Giovanni Ciriello. Cellcharter reveals spatial cell niches associated with tissue remodeling and cell plasticity. Nature genetics, 56(1):74–84, 2024.

[19] Yue You, Yuting Fu, Lanxiang Li, Zhongmin Zhang, Shikai Jia, Shihong Lu, Wenle Ren, Yifang Liu, Yang Xu, Xiaojing Liu, et al. Systematic comparison of sequencing-based spatial transcriptomic methods. Nature methods, 21(9):1743–1754, 2024.

[20] Jinyun Niu, Donghai Fang, Jinyu Chen, Yi Xiong, Juan Liu, and Wenwen Min. Spabatch: Deep learning-based cross-slice integration and 3d spatial domain identification in spatial transcriptomics. Advanced Science, n/a(n/a):e09090. doi: 10.1002/advs.202509090. URL https://advanced.onlinelibrary.wiley.com/doi/abs/10.1002/advs.202509090.

[21] Zhiyuan Yuan, Fangyuan Zhao, Senlin Lin, Yu Zhao, Jianhua Yao, Yan Cui, Xiao-Yong Zhang, and Yi Zhao. Benchmarking spatial clustering methods with spatially resolved transcriptomics data. Nature Methods, 21(4):712–722, 2024.

[22] Liyang Song, Wenhao Chen, Junren Hou, Minmin Guo, and Jian Yang. Spatially resolved mapping of cells associated with human complex traits. Nature, 641(8064):932–941, 2025.

[23] Hongen Kang, Xiaoxi Jing, Jiecong Lin, Siyu Pan, Jiaxiang Zhang, Junpeng Zhao, and Yajie Zhao. Spatial gwas atlas: a knowledgebase for decoding the genetic architecture of complex traits in spatial resolution. Nucleic Acids Research, 54(D1):D1301–D1308, 2026.

[24] Xi Hu, Aoqi Wang, Huan Yu, Pora Kim, and Xiaobo Zhou. Spatial2gwas: a database for linking spatial transcriptomic regions with gwas traits. Nucleic Acids Research, 54(D1):D1309–D1320, 2026.

[25] Amanda Janesick, Robert Shelansky, Andrew D Gottscho, Florian Wagner, Stephen R Williams, Morgane Rouault, Ghezal BeliakoK, Carolyn A Morrison, Michelli F Oliveira, Jordan T Sicherman, et al. High resolution mapping of the tumor microenvironment using integrated single-cell, spatial and in situ analysis. Nature communications, 14(1):8353, 2023.

[26] Sang Uk Han, Tae Hee Kwak, Kyu Hee Her, Young-Hun Cho, Chul-won Choi, Ho Jae Lee, Suntaek Hong, Yoon Soo Park, YS Kim, Tae Aug Kim, et al. Ceacam5 and ceacam6 are major target genes for smad3-mediated tgf-*β* signaling. Oncogene, 27(5):675–683, 2008.

[27] JeKrey R Moffitt, Dhananjay Bambah-Mukku, Stephen W Eichhorn, Eric Vaughn, Karthik Shekhar, Julio D Perez, Nimrod D Rubinstein, Junjie Hao, Aviv Regev, Catherine Dulac, et al. Molecular, spatial, and functional single-cell profiling of the hypothalamic preoptic region. Science, 362 (6416):eaau5324, 2018.

[28] Shun Wang, Baozhen Yao, Haiju Zhang, Liping Xia, Shiqian Yu, Xia Peng, Dan Xiang, and Zhongchun Liu. Comorbidity of epilepsy and attention-deficit/hyperactivity disorder: a systematic review and meta-analysis. Journal of Neurology, 270(9):4201–4213, 2023.

[29] Hu Zeng, Jiahao Huang, Haowen Zhou, William J Meilandt, Borislav Dejanovic, Yiming Zhou, Christopher J Bohlen, Seung-Hye Lee, Jingyi Ren, Albert Liu, et al. Integrative in situ mapping of single-cell transcriptional states and tissue histopathology in a mouse model of alzheimer’s disease. Nature neuroscience, 26(3):430–446, 2023.

[30] Naomi Habib, Cristin McCabe, Sedi Medina, Miriam Varshavsky, Daniel Kitsberg, Raz Dvir-Szternfeld, Gilad Green, Danielle Dionne, Lan Nguyen, Jamie L Marshall, et al. Disease-associated astrocytes in alzheimer’s disease and aging. Nature neuroscience, 23(6):701–706, 2020.

[31] Hansruedi Mathys, Jose Davila-Velderrain, Zhuyu Peng, Fan Gao, Shahin Mohammadi, Jennie Z Young, Madhvi Menon, Liang He, Fatema Abdurrob, Xueqiao Jiang, et al. Single-cell transcriptomic analysis of alzheimer’s disease. Nature, 570(7761):332–337, 2019.

[32] Cui Zhang, Hao Qi, Dongjing Jia, Jingting Zhao, Chengyuan Xu, Jing Liu, Yangfeng Cui, Jiajian Zhang, Minzhe Wang, Ming Chen, et al. Cognitive impairment in alzheimer’s disease fad4t mouse model: Synaptic loss facilitated by activated microglia via c1qa. Life Sciences, 340: 122457, 2024.

[33] Borislav Dejanovic, TiKany Wu, Ming-Chi Tsai, David Graykowski, Vineela D Gandham, Christopher M Rose, Corey E Bakalarski, Hai Ngu, Yuanyuan Wang, Shristi Pandey, et al. Complement c1q-dependent excitatory and inhibitory synapse elimination by astrocytes and microglia in alzheimer’s disease mouse models. Nature Aging, 2(9):837–850, 2022.

[34] Yuhong Li, Xiaofan Zhang, and Deming Chen. Csrnet: Dilated convolutional neural networks for understanding the highly congested scenes. In Proceedings of the IEEE conference on computer vision and pattern recognition, pages 1091–1100, 2018.

[35] Herbert Robbins and Sutton Monro. A stochastic approximation method. The annals of mathematical statistics, pages 400–407, 1951.

[36] Léon Bottou, Frank E Curtis, and Jorge Nocedal. Optimization methods for large-scale machine learning. SIAM review, 60(2):223–311, 2018.

[37] Hanqing Zeng, Hongkuan Zhou, Ajitesh Srivastava, Rajgopal Kannan, and Viktor Prasanna. Graphsaint: Graph sampling based inductive learning method. arXiv preprint arXiv:1907.04931, 2019.

[38] Petar Veličković, Guillem Cucurull, Arantxa Casanova, Adriana Romero, Pietro Liò, and Yoshua Bengio. Graph attention networks. In International Conference on Learning Representations, page n/a, 2018.

[39] Xiaotong Li, Yongxing Dai, Yixiao Ge, Jun Liu, Ying Shan, and Lingyu Duan. Uncertainty modeling for out-of-distribution generalization. In International Conference on Learning Representations, page n/a, 2022.

[40] Yue Liu, Ke Liang, Jun Xia, Sihang Zhou, Xihong Yang, Xinwang Liu, and Stan Z Li. Dink-net: Neural clustering on large graphs. In International Conference on Machine Learning, pages 21794–21812. PMLR, 2023.

[41] Aaron van den Oord, Yazhe Li, and Oriol Vinyals. Representation learning with contrastive predictive coding. arXiv preprint arXiv:1807.03748, 2018.

[42] Wei Liu, Bo Wang, Yuting Bai, Xiao Liang, Li Xue, and Jiawei Luo. Spagic: graph-informed clustering in spatial transcriptomics via self-supervised contrastive learning. Briefings in Bioinformatics, 25(6):bbae578, 11 2024. ISSN 1477-4054. doi: 10.1093/bib/bbae578. URL 10.1093/bib/bbae578.

[43] Ilya Loshchilov and Frank Hutter. Decoupled weight decay regularization. In International Conference on Learning Representations, page n/a, 2019.

[44] Matthias Fey and Jan Eric Lenssen. Fast graph representation learning with pytorch geometric. arXiv preprint arXiv:1903.02428, 2019.

[45] Luca Scrucca, Michael Fop, T Brendan Murphy, and Adrian E Raftery. mclust 5: clustering, classification and density estimation using gaussian finite mixture models. The R journal, 8(1): 289, 2016.

[46] Vincent A Traag, Ludo Waltman, and Nees Jan Van Eck. From louvain to leiden: guaranteeing well-connected communities. Scientific reports, 9(1):1–12, 2019.

[47] Pablo A. Estévez, Michel Tesmer, Claudio A. Perez, and Jacek M. Zurada. Normalized mutual information feature selection. IEEE Trans. Neural Networks, 20(2):189–201, 2009. doi: 10.1109/TNN.2008.2005601. URL 10.1109/TNN.2008.2005601.

[48] Rochelle L Castillo, Ikjot Sidhu, Igor Dolgalev, Tinyi Chu, Aleksandr Prystupa, Ipsita Subudhi, Di Yan, Piotr Konieczny, Brandon Hsieh, Rebecca H Haberman, et al. Spatial transcriptomics stratifies psoriatic disease severity by emergent cellular ecosystems. Science immunology, 8(84): eabq7991, 2023.

[49] DM Calcagno, N Taghdiri, VK Ninh, JM Mesfin, A Toomu, R Sehgal, J Lee, Y Liang, JM Duran, E Adler, et al. Single-cell and spatial transcriptomics of the infarcted heart define the dynamic onset of the border zone in response to mechanical destabilization. Nature cardiovascular research, 1(11):1039–1055, 2022.

[50] Karel van Duijvenboden, Dennis EM de Bakker, Joyce CK Man, Rob Janssen, Marie Günthel, Matthew C Hill, Ingeborg B Hooijkaas, Ingeborg van der Made, Petra H van der Kraak, Aryan Vink, et al. Conserved nppb+ border zone switches from mef2-to ap-1–driven gene program. Circulation, 140(10):864–879, 2019.

[51] Upendra Chalise, Michael J Daseke, William J Kalusche, Shelby R Konfrst, Jocelyn R Rodriguez-Paar, Elizabeth R Flynn, Leah M Cook, Mediha Becirovic-Agic, and Merry L Lindsey. Macrophages secrete murinoglobulin-1 and galectin-3 to regulate neutrophil degranulation after myocardial infarction. Molecular Omics, 18(3):186–195, 2022.

[52] Mengting Chen, Yaling Wang, Mei Wang, San Xu, Zixin Tan, Yisheng Cai, Xin Xiao, Ben Wang, Zhili Deng, and Ji Li. Keratin 6a promotes skin inflammation through jak1-stat3 activation in keratinocytes. Journal of Biomedical Science, 32(1):47, 2025.

[53] Bruno Marcel Silva de Melo, Flavio Protasio Veras, Pascale Zwicky, Diogenes Lima, Florian Ingelfinger, Timna Varela Martins, Douglas da Silva Prado, Stefanie Schärli, Gabriel Publio, Carlos Hiroji Hiroki, et al. S100a9 drives the chronification of psoriasiform inflammation by inducing il-23/type 3 immunity. Journal of Investigative Dermatology, 143(9):1678–1688, 2023.

