## Supplementary material for "Mapping disease traits onto spatiotemporal domain landscapes through scalable multi-sample integration of spatial transcriptomics": Supplmental files

### S1. Supplementary Figures

**Unified ablation study of STUltra.** We conducted a unified ablation study to evaluate the contribution of each component of STUltra. The results are shown in Figure S1. Using paired human colorectal cancer Visium HD sections, we conducted a systematic sensitivity analysis of three key settings in the STUltra integration pipeline under fixed clustering resolution and training budget: the spatial neighborhood size (KNN), the cross-batch alignment strength (loss weight $\alpha$), and the number of interval-sampled batches used (`num_batch`). The evaluation jointly considered whether histological clusters remained clearly organized, whether clusters were well separated, and whether samples from different batches or sections could still be distinguished or had been effectively mixed in the embedding space, thereby characterizing how the model balances preservation of biological structure against removal of technical batch effects.

Overall, the neighborhood size had a relatively mild influence on the results, suggesting that within the tested range, local graph construction was not the primary bottleneck for integration performance. In contrast, the alignment strength $\alpha$ dominated the performance trade-off. Under weaker alignment, cells were more likely to form structurally coherent groups according to histological states, although noticeable batch effects remained. Increasing the alignment strength substantially reduced these biological structures while improving batch mixing, at the cost of lower clustering quality. Increasing `num_batch` further strengthened the debatching tendency and improved batch mixing metrics, but simultaneously reduced cluster coherence and spatial continuity. When the interval-sampled batch partition became excessively fine-grained, the integrated representations could even become fragmented.

Therefore, hyperparameter selection in STUltra fundamentally reflects a trade-off between tissue structure fidelity and cross-batch comparability. If the primary goal is to identify disease-associated cell states or spatial domains, lighter alignment settings are generally preferable. In contrast, if the objective is to construct a unified atlas directly comparable across sections or batches, stronger alignment ($\alpha$) or larger `num_batch` values may be beneficial, although this comes with the risk of blurred biological clustering. The current default configuration represents a compromise between these two objectives and is better viewed as a practical starting point for downstream analysis rather than a task-specific optimum.

**Resolution parameter in Louvain clustering.** The resolution parameter in the Louvain algorithm determines the granularity of the detected communities and directly influences the number of clusters obtained. A lower resolution yields fewer, larger clusters, while a higher resolution produces more, smaller clusters. In our experiments, we select the resolution parameter by considering both the cellular density of each dataset and the parameter settings recommended in previous ST studies. For datasets with low spatial resolution or sparse spatial input unit distributions, we set the resolution in the range of 0.2 – 0.5 to maintain biologically meaningful groupings without fragmenting tissue regions. In contrast, for high-resolution ST slices, we increase the resolution to 0.4–1.0 to preserve local transcriptional heterogeneity. This adaptive choice ensures that the clustering scale is consistent with the intrinsic resolution and biological complexity of each dataset.

**Cardiac ischemic injury case study.** We examined whether STUltra could capture dynamic spatial reorganization during disease progression, using a mouse cardiac ischemic injury dataset comprising four ST slices taken at key post-injury time points (1 hour, 4 hours, 1 day, 7 days). The spatial domains include border zones (BZ1 and BZ2), infarct zones (IZ), and remote zones (RZ). BZ1 and BZ2 represent transitional regions between injured and healthy tissue, IZ corresponds to necrotic tissue, and RZ denotes uninjured myocardium. STUltra accurately identified these domains and

| | | Cluster | | | Batch | | |
|---|---|---|---|---|---|---|---|
| **Ablation** | Value | Silhouette | Calinski-H | Davies-B | Sil. batch | iLISI | Graph conn. |
| **KNN** | 10 | 0.13 | 402.2 | 1.92 | 0.020 | 1.48 | 0.73 |
| | 20 | 0.13 | 393.9 | 1.88 | 0.016 | 1.49 | 0.97 |
| | 50 | 0.13 | 397.6 | 1.98 | 0.012 | 1.51 | 0.92 |
| | 100 | 0.13 | 394.9 | 1.96 | 0.011 | 1.52 | 0.92 |
| **Alpha** | 0.2 | 0.18 | 827.7 | 1.65 | 0.025 | 1.43 | 0.93 |
| | 0.5 | 0.18 | 732.8 | 1.68 | 0.019 | 1.44 | 0.94 |
| | 1 | 0.16 | 548.5 | 1.76 | 0.014 | 1.46 | 0.88 |
| | 2 | 0.15 | 510.8 | 1.83 | 0.015 | 1.47 | 0.85 |
| | 4 | 0.14 | 441.5 | 1.86 | 0.011 | 1.51 | 0.83 |
| | 5 | 0.13 | 407.4 | 1.90 | 0.012 | 1.52 | 0.67 |
| **Num batch** | 2 | 0.13 | 407.2 | 1.92 | 0.011 | 1.52 | 0.85 |
| | 3 | 0.10 | 351.7 | 2.05 | 0.008 | 1.54 | 0.65 |
| | 4 | 0.11 | 369.5 | 2.03 | 0.007 | 1.55 | 0.78 |
| | 5 | 0.12 | 366.5 | 2.00 | 0.005 | 1.58 | 0.39 |

Figure S1 | Unified ablation study of STUltra.

revealed distinct spatial localization compared with manual annotations (Fig. S3a); for example, BZ1 was concentrated near the vascular wall at early stages, whereas the annotation appeared diffusely distributed within the vessel. Spatial expression of injury-responsive markers confirmed domain specificity of our results (Fig. S3b). *Nppb* is a marker of cardiac stress [50], and *Lgals3* is associated with macrophage infiltration [51]. Both genes were localized to BZ regions, consistent with their roles in early injury response and tissue repair. We further performed pseudotime trajectory analysis on STUltra's integrated embeddings, which revealed a continuous progression of molecular states across the injury time points (Fig. S3c,d, Fig. S16). This analysis disclosed a mechanism of dynamic transitions between tissue regions following cardiac injury, providing insights into the temporal ordering of cellular responses. For example, when the expression of RZ marker genes (*Eno3*, *Tcap*) was gradually reduced in magnitude, BZ1 marker genes (*Nppb*, *Cilp*) and BZ2 marker genes (*Fn1*, *Xirp2*) were sequentially activated. These sequential activations reflect the spatial and temporal remodeling of the myocardium during injury, including the activation of fibroblasts and stress-responsive cardiomyocytes.

**Psoriatic skin case study.** We assessed whether STUltra could directly distinguish pathological from healthy tissue architecture in a chronic inflammatory disease. Psoriasis, one of the most common immune-mediated diseases, provides an ideal model for such comparison through pair-matched lesional and contralateral non-lesional skin samples [48]. We detected 15 clusters (Fig. S4a), and found that Clusters 5 and 6 were markedly expanded in the lesional skin in comparison with the non-lesional skin (Fig. S4b). These two clusters were spatially located near the dermal layer (Cluster 3) and were also closely grouped in the UMAP space, suggesting a pathological association feature

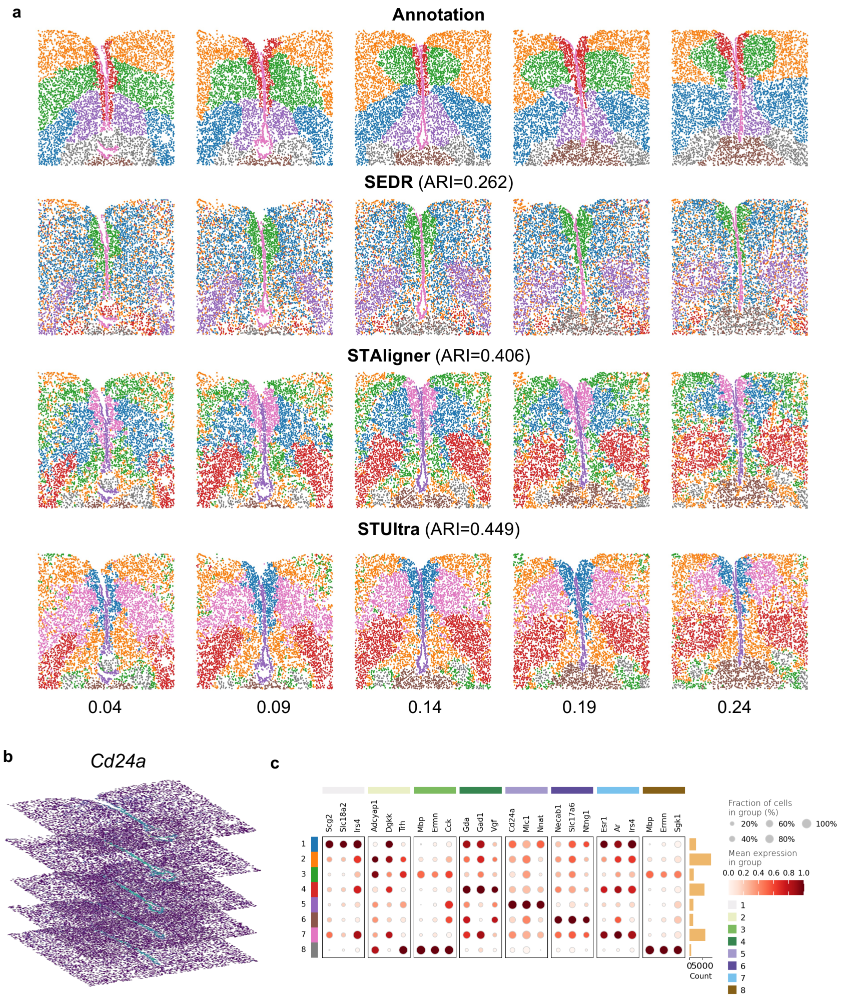


Figure S2 | Analysis on five sequential slices of the mouse hypothalamic preoptic area, profiled by the MERFISH platform. (a) Spatial domains identified using STUltra and other methods. (b) Dot plot of differential analysis across eight clusters. (c) STUltra performed a 3D spatial reconstruction of the marker gene *Cd24a* within the V3 region (Domain 5).

of psoriasis. Marker gene analysis for these subpopulations, together with their spatial expression patterns, confirmed a local up-regulation and strong spatial co-localization of psoriasis-associated differentially expressed genes (Fig. S4c,e). For example, *KRT6A* was recognized as a marker of hyperproliferative keratinocytes [52], while *S100A9* has been reported to mediate the chronification of inflammation [53]. These differentially expressed genes were enriched in multiple Gene Ontology (GO) terms, including apoptotic process and T cell receptor signaling (Fig. S4d), reflecting the key molecular pathways that were active in the psoriatic lesions.

#### S2. Supplementary tables

Table S1 | Summary of the ST data used in this study.

| Tissue | Platform | Slice id | # of input units | Related figures |
|---|---|---|---|---|
| DLPFC | 10x Visium | 151507 | 4,226 | Fig. 2, Fig. S7-S6 |
| | | 151508 | 4,384 | |
| | | 151509 | 4,789 | |
| | | 151510 | 4,634 | |
| Mouse hypothalamus | MERFISH | 0.04 | 5557 | Fig. S2 |
| | | 0.09 | 5,926 | |
| | | 0.14 | 5,803 | |
| | | 0.19 | 5,803 | |
| | | 0.24 | 5,543 | |
| Mouse brain | 10x Visium | Sagittal-anterior | 2,695 | Fig. 2 |
| | | Sagittal-posterior | 3,355 | |
| Mouse brain | Visium HD | brain_visium_local_hd | 579,500 | Fig. 5 |
| | BMK S1000 | brain_bmk_1 | 249,130 | |
| | Stereo-Seq v2 | brain_stomic | 577,094 | |
| Mouse embryos | Stereo-seq | E9.5_E1S1 | 5913 | Fig. 4, Fig. S10-S11 |
| | | E10.5_E2S1 | 8,494 | |
| | | E11.5_E1S1 | 30,124 | |
| | | E12.5_E1S1 | 51,365 | |
| | | E13.5_E1S1 | 77,369 | |
| | | E14.5_E1S1 | 102,519 | |
| | | E15.5_E1S1 | 113,350 | |
| | | E16.5_E1S1 | 121,767 | |
| Human colorectal cancer | Visium HD | P1 | 507,684 | Fig. 3, Fig. S12-S14 |
| | | P2 | 545,913 | |
| Human breast cancer | Xenium | Rep1 | 167,782 | Fig. S8, S15 |
| | | Rep2 | 118,708 | |
| Human skin | 10x Visium | ST_21_NL | 955 | Fig. S4 |
| | | ST_22_L | 654 | |
| Mouse Heart | 10x Visium | 1HR MI | 2,468 | Fig. S3, S16 |
| | | 4HR MI | 2,578 | |
| | | Day3 MI biol rep 1 | 2,236 | |
| | | Day7 MI biol rep 1 | 2,313 | |
| Alzheimer's disease | STARmap PLUS | 8months-control | 8,506 | Fig. 6 |
| | | 8months-disease | 8,186 | |
| | | 13months-control | 8,034 | |
| | | 13months-disease | 1,0372 | |

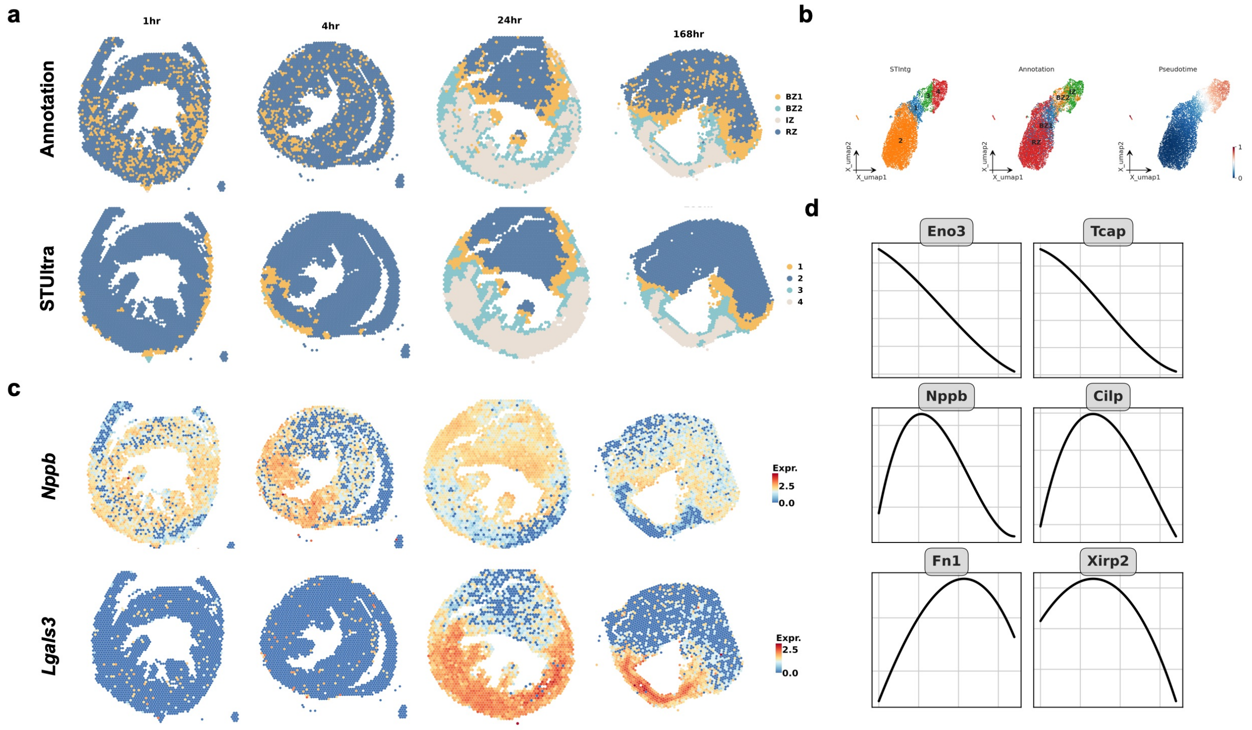


Figure S3 | **STUltra identifies disease-associated dynamics on ST slices in four mouse heart slices from an ischemic injury dataset, covering four time points (1 hour, 4 hours, 1 day, and 7 days).** (a) Manual annotations and spatial domains identified by STUltra. The spatial regions include border zone 1/2 (BZ1, BZ2), infarct zones (IZ), and remote zones (RZ). (b) Spatial expression maps of marker genes (*Nppb*, *Lgals3*) in border zones. (c) UMAP plots of STUltra embeddings colored by spatial domain, manual annotation, and pseudotime. (d) Trajectory analysis of marker genes along the direction of pseudotime.

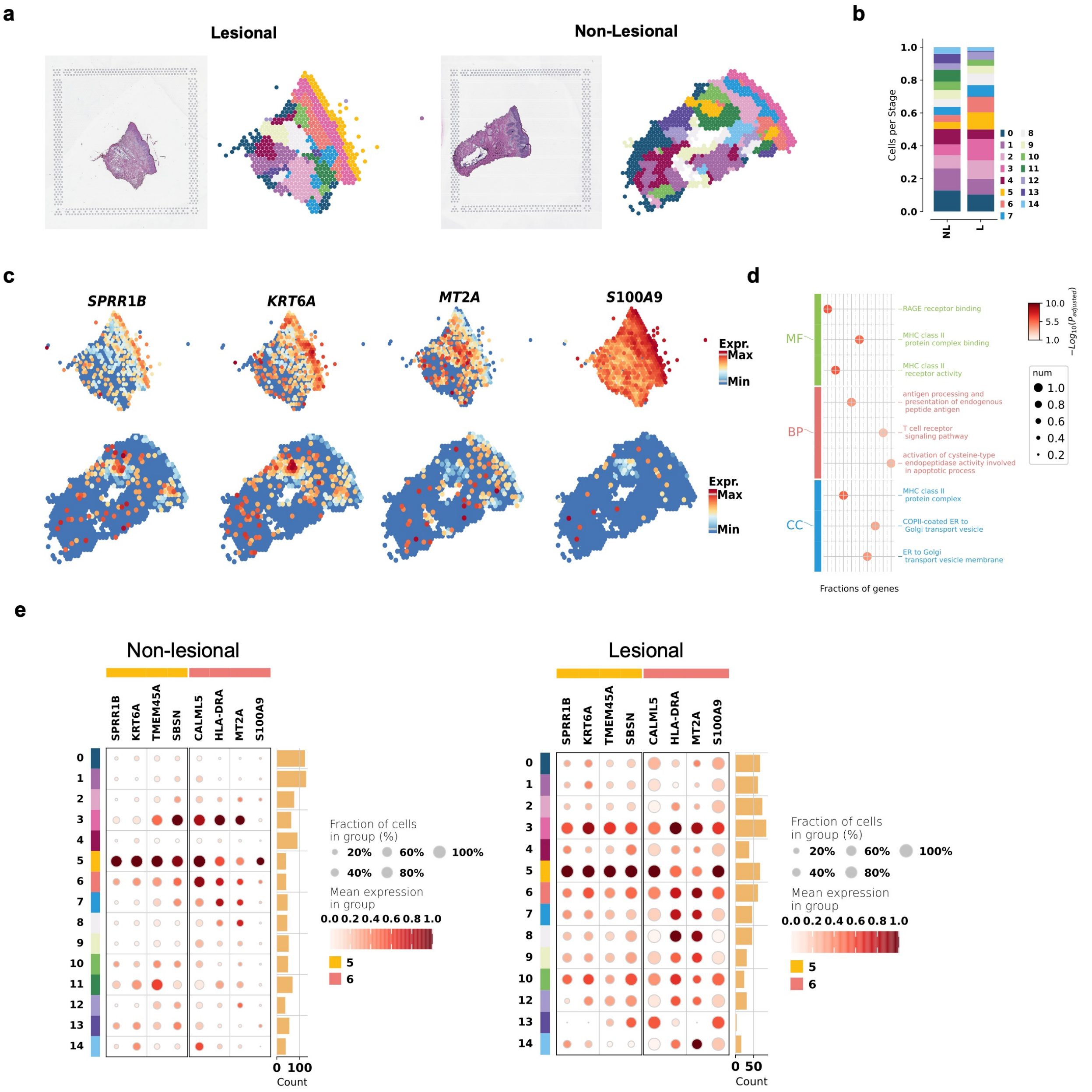


Figure S4 | (a) H&E-stained images and visualization of aligned spatial domains identified by STUltra on human psoriatic lesional and non-lesional skin slices. (b) Stacked bar plot of cell-type proportions across spatial domains. (c) Spatial expression maps of four marker genes (*SPRR1B*, *KRT6A*, *MT2A*, *S100A9*) for Cluster 6 in human psoriatic lesional and non-lesional skin slices. (d) Gene Ontology (GO) analysis comparing Cluster 5 with other clusters. (e) The dot plot of the highly expressed genes in STUltra's clusters in human psoriatic dataset.

**Algorithm 1** STUltra algorithm.

**Require:** $\{\mathbf{X}_s\}_{s=1}^{S}$: gene expression matrices of $S$ slices; $\{\text{Coord}_s\}_{s=1}^{S}$: spatial coordinates; $K$: number of highly variable genes; $\theta_{\text{graph}} \in \{k, r\}$: graph parameter ($k$-NN or radius $r$); $T_1$, $T_2$: iteration limits
**Ensure:** $\mathbf{Z}_{\text{final}}$: integrated latent embeddings

```
Preprocessing:
for each slice s do
    Normalize X_s and select Top-K HVGs;
    Partition slice into subgraphs {G^(m)_sub,s} by interval δ;
    Build adjacency {A^(m)_sub,s} via θ_graph;
end for
Compute shared HVGs across slices and update {X_s};
For each index m, group {G^(m)_sub,s}_{s=1}^S into batch G^(m)_B;
Collect all batches 𝒢_B = {G^(m)_B} as the training set;

Stage 1: Robust GAE Pre-training
Initialize encoder and decoder weights
while not converged and t_1 < T_1 do
    for each 𝒢_B do
        Training Encoder and decoder via Eq. 3–5;
        Apply uncertainty estimation via Eq. 7–9;
        Compute ℒ_rec, ℒ_dis via Eq. 6–11;
        Update parameters with ℒ_1 via Eq. 12;
    end for
end while
Initialize Z_2 = Z_1;

Stage 2: Batch-Disentangled Contrastive Learning
Construct initial MNN-based contrastive pairs;
while not converged and t_2 < T_2 do
    if t_2 % 100 == 0 or t_2 == 0 then
        Recompute MNN pairs using current embeddings Z_2;
    end if
    for each 𝒢_B do
        Update Encoder with ℒ_2 and MNN pairs via Eq. 13–14;
        Compute Z_2 via Encoder;
    end for
    t_2 ← t_2 + 1;
end while
return Z_final ← Z_2.
```

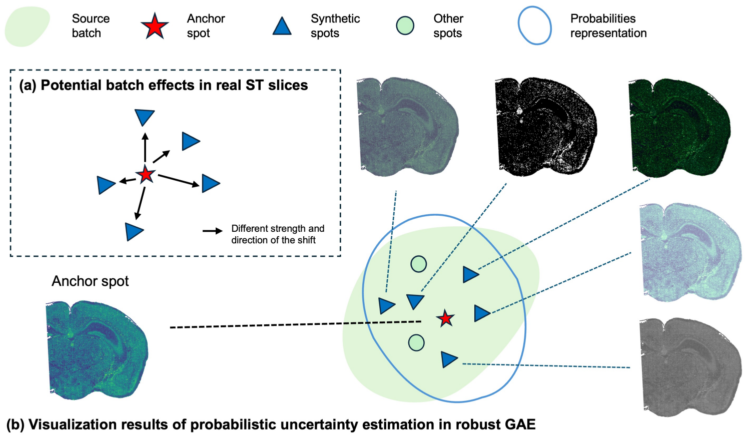


Figure S5 | Overview of probabilistic uncertainty estimation. (a) Potential batch effects in real ST slices. (b) Visualization results of probabilistic uncertainty estimation in robust graph autoencoder.

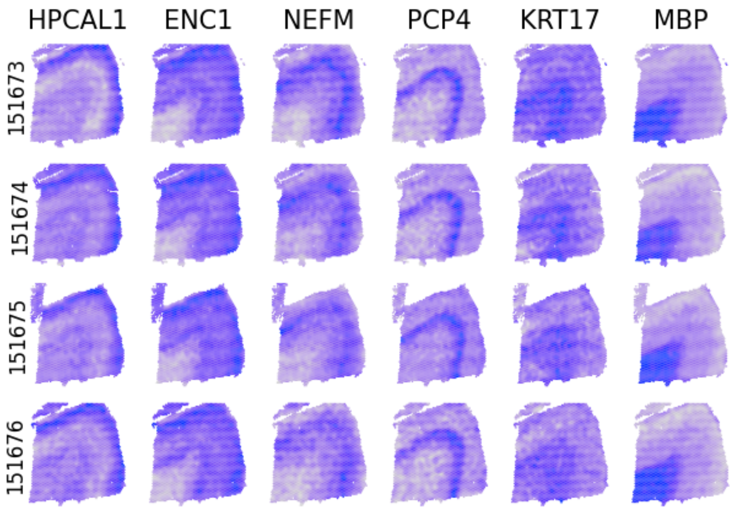


Figure S6 | The spatial expression of the layer 1-6 and WM marker gene in DLPFC dataset.

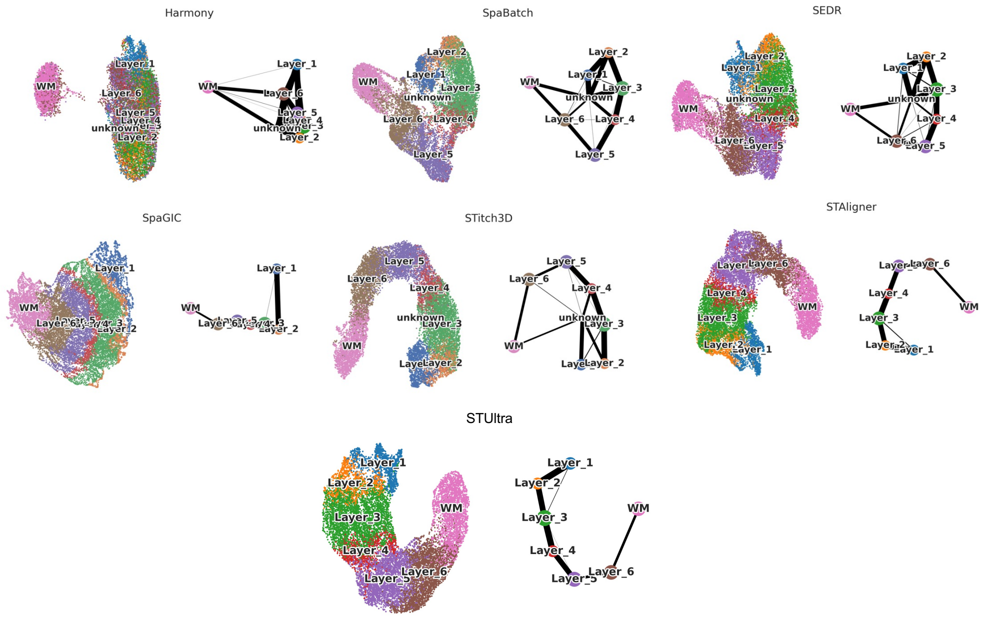


Figure S7 | The UMAP visualization and domain trajectory was compared visually using PAGA.

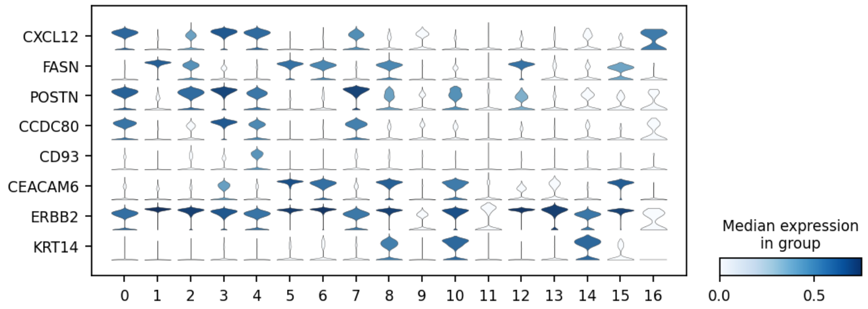


Figure S8 | The violin plot of the highly expressed genes in STUltra's clusters of the human breast cancer(BC) 10x Xenium dataset.

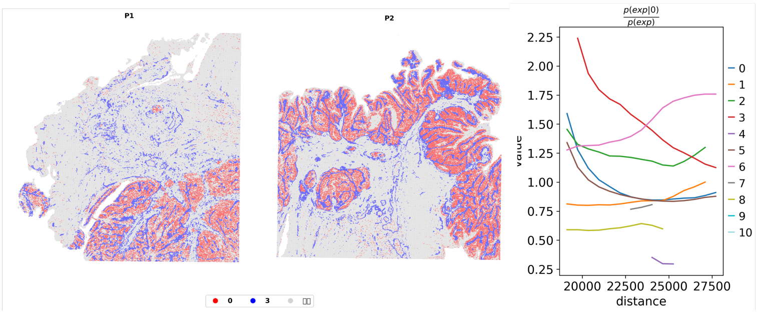


Figure S9 | Visualization of the tumor periphery in Visium HD human colorectal cancer slices. Tumor-cell bins are shown in red, and bins located within 50$\mu$m from the tumor boundary are highlighted in blue. Spatial co-occurrence of different cell types relative to Cluster 0 (tumor), computed using Squidpy. Cluster 3 (fibroblasts) is closest to the tumor edge.

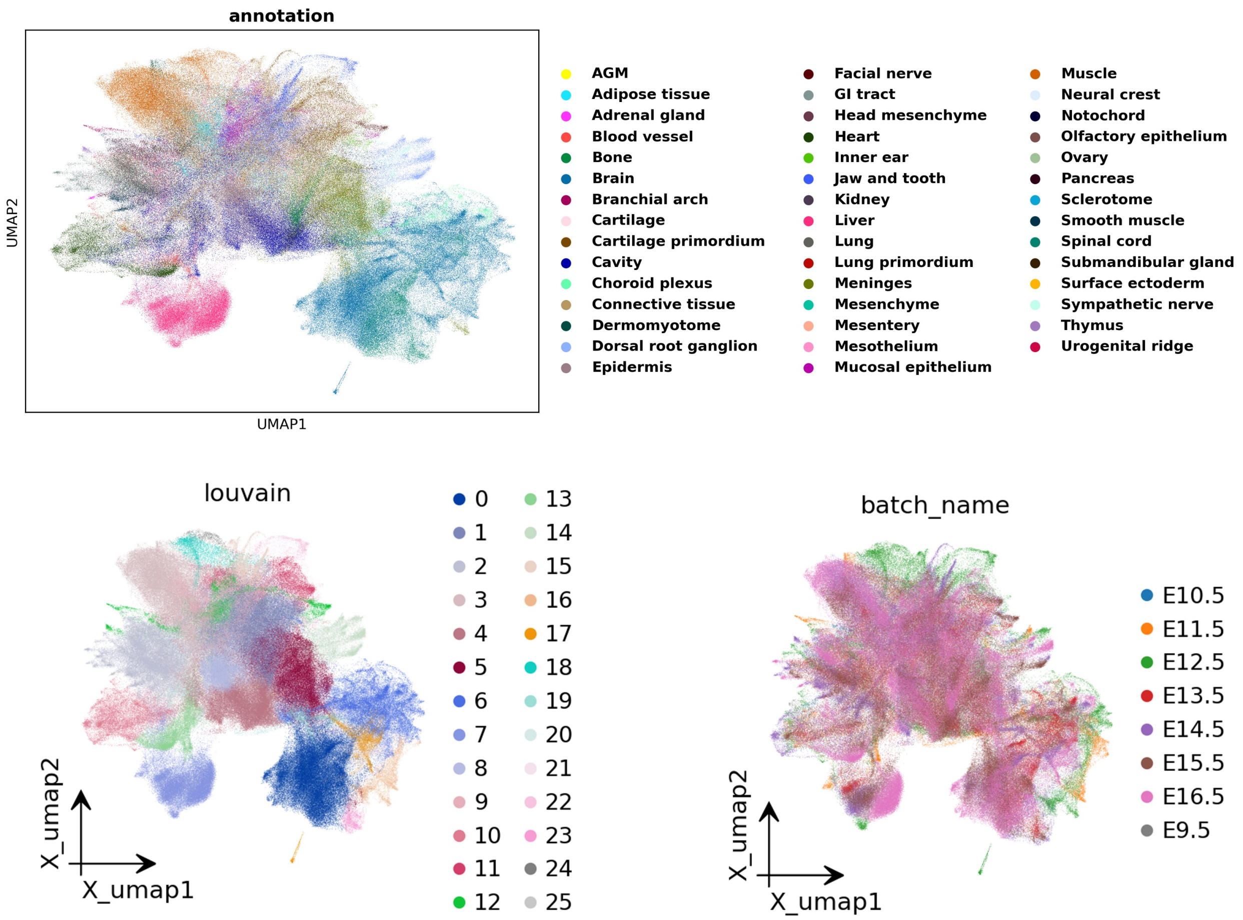


Figure S10 | The UMAP visualization colored by annotation, batch, and STUltra in mouse embryos dataset.

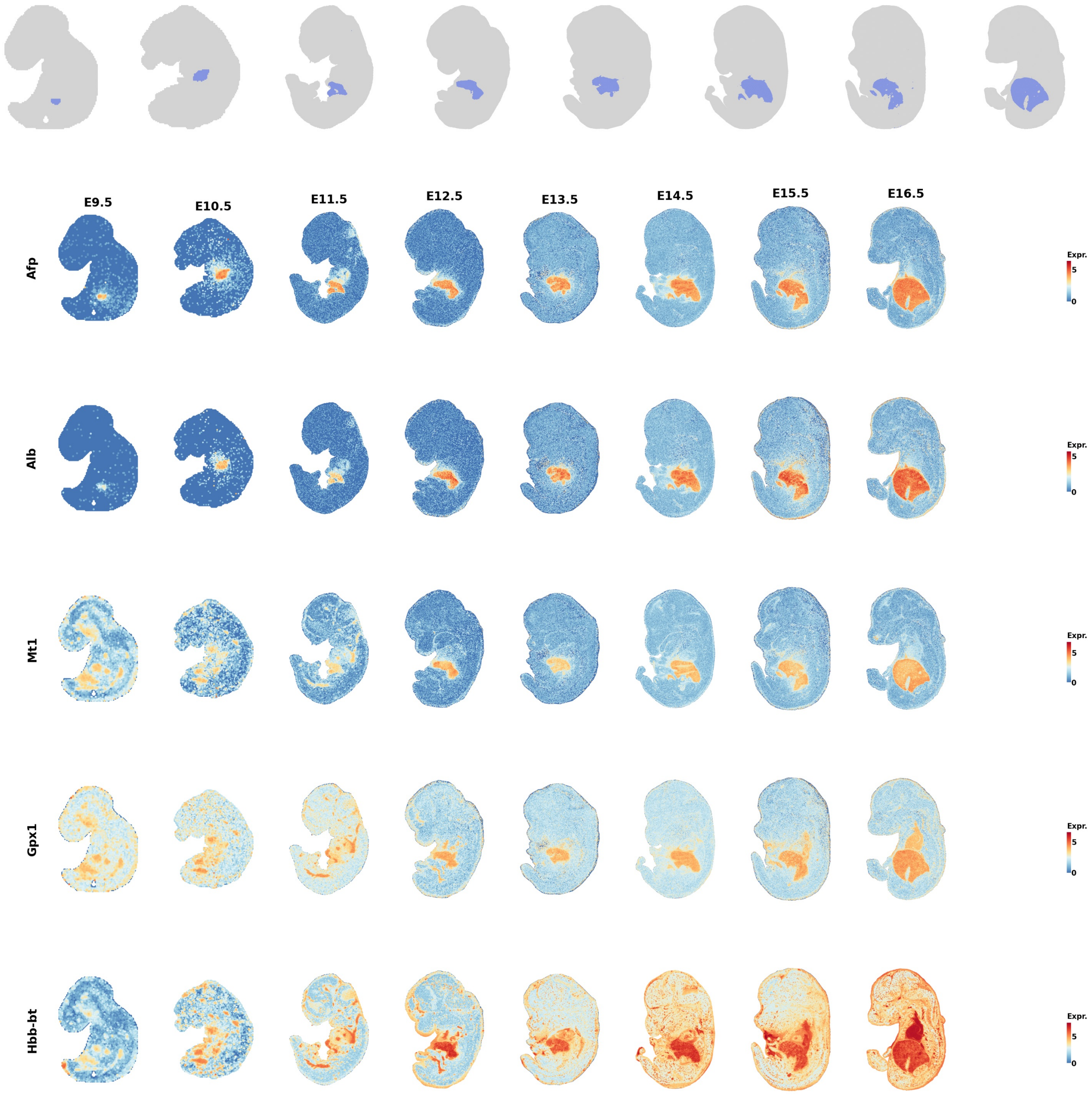


Figure S11 | The liver and spatial expressions returned by STUltra in mouse embryos dataset.

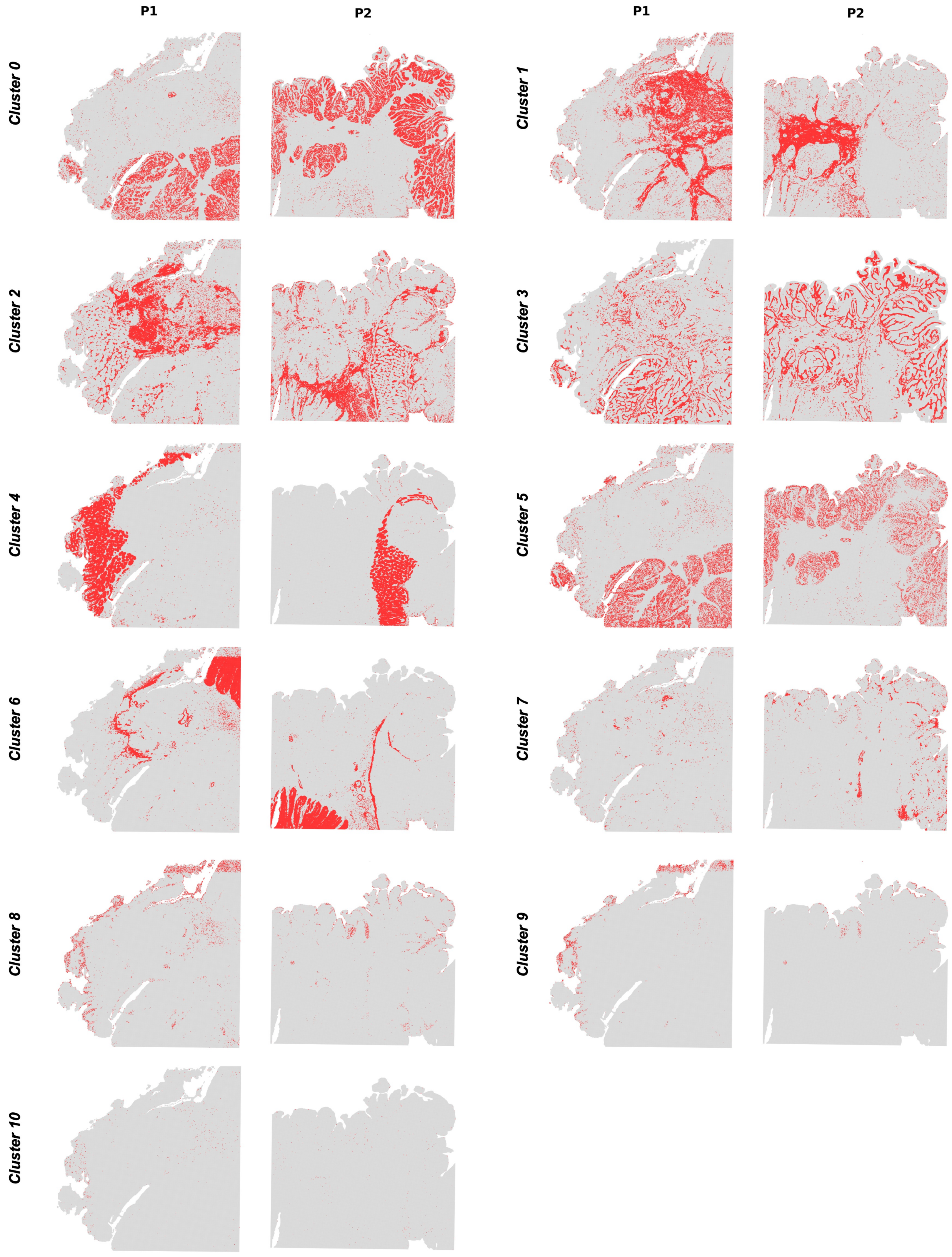


Figure S12 | The spatial visualization returned by STUltra in CRC dataset.

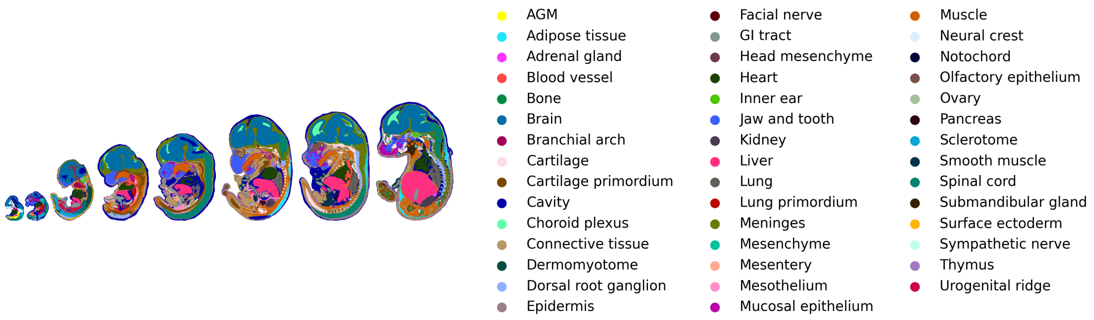


Figure S13 | The spatial domains returned by annotation in mouse embryos dataset.

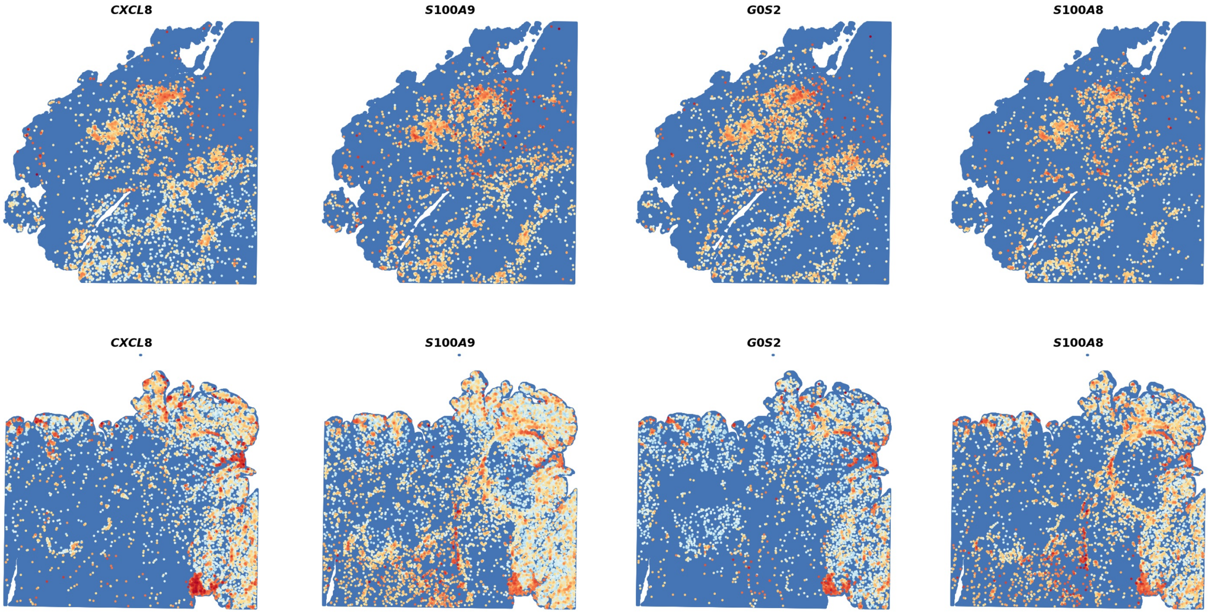


Figure S14 | The Cluster 7 and spatial expressions returned by STUltra in CRC dataset.

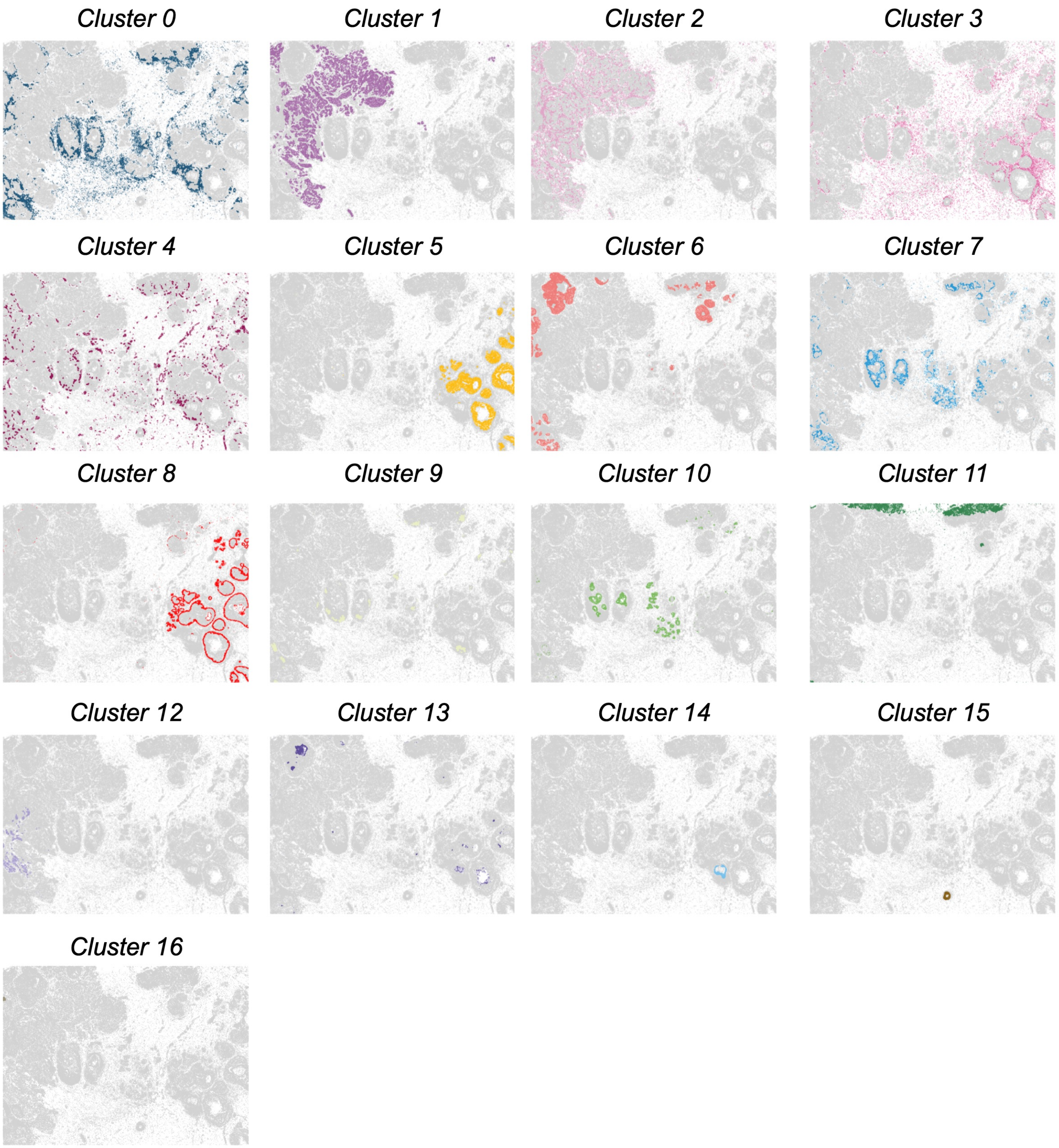


Figure S15 | The spatial visualization separately returned by STUltra in the slice Rep1 of human BC 10x Xenium dataset. STUltra effectively distinguishes and localizes four spatially distinct tumor domains (cluster 1, 2, 5, and 8).

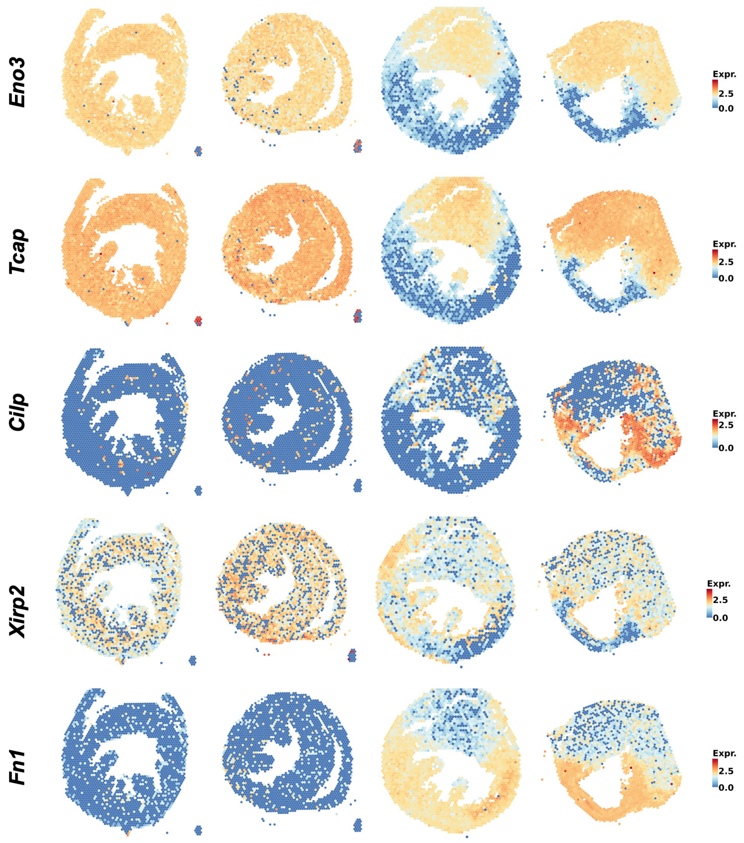


Figure S16 | The spatial expressions returned by STUltra in mouse heart dataset.
